# Re-direction of phagosomes to the recycling expulsion pathway by a fungal pathogen

**DOI:** 10.1101/2022.05.18.492126

**Authors:** Lei-Jie Jia, Muhammad Rafiq, Lukáš Radosa, Peter Hortschansky, Cristina Cunha, Zoltán Cseresnyés, Thomas Krüger, Franziska Schmidt, Thorsten Heinekamp, Maria Straßburger, Bettina Löffler, Torsten Doenst, João F. Lacerda, António Campos, Marc Thilo Figge, Agostinho Carvalho, Olaf Kniemeyer, Axel A. Brakhage

**Author notes:** Correspondence (A.A.B).

## Abstract

The analysis of host-pathogen interactions bears the potential to discover novel pathogenicity mechanisms and to obtain novel insights into basic mechanisms of cell biology. Here, we obtained unprecedented insight into both. We discovered that the HscA protein on the conidial surface of the clinically important human-pathogenic fungus *Aspergillus fumigatus* acts as an effector protein. It inhibits phagosome maturation and reprograms phagosomes for expulsion of conidia. HscA anchors the human p11 protein to phagosomes. p11 is a decisive factor for targeting phagosomes either to the degradative or secretory pathway. The relevance of our findings is indicated by the identification of an SNP in the non-coding region of the human p11 gene that affects its translation and is associated with heightened susceptibility to invasive pulmonary aspergillosis.

## Introduction

A basic question of cell biology is the molecular understanding of the sorting of internalized cargoes, which includes the decision whether endosomes and their cargo enter the degradative or non-degradative, recycling pathways (Cullen and Steinberg, 2018; Pauwels et al., 2017). In the non-degradative pathways, cargos are either redirected to the cell surface, secreted to the environment, maintained in intracellular vesicles, or even transferred to other cells (Brakhage et al., 2021; Cullen and Steinberg, 2018; Kinchen and Ravichandran, 2008; Serra and Sundaram, 2021). Erroneous decisions in endosomal sorting are associated with various diseases, including neurological and immunological disorders, as well as cancer (Yarwood et al., 2020). The investigation of pathogens that interfere with phagosome maturation in cells is important to understand pathogenicity but also helps to identify host proteins controlling the fate of endosomes (Ledvina et al., 2018; Schmidt et al., 2020; Walpole et al., 2020; Xie et al., 2021). Until now, there is only limited knowledge about such proteins that are decisive for endosomes to enter the degradative or secretory pathways (van Niel et al., 2018; Wei et al., 2020).

Here, by analyzing the intracellular processing of spores of the medically important pathogenic fungus *Aspergillus fumigatus* we identified a novel effector molecule and a regulatory node controlling the fate of endosomes. *A. fumigatus* is an opportunistic human pathogen, which causes disseminated infections in immunocompromised patients (Brakhage, 2005; Dagenais and Keller, 2009; Kousha et al., 2011; Latgé and Chamilos, 2019; Taccone et al., 2015). The fungus produces conidia, asexually produced spores, that are released into the air and are continuously inhaled. Because of their small size of 2–3 µm, conidia can easily reach the lung alveoli (Brakhage and Langfelder, 2002). Without an effective immune response, inhaled conidia germinate and grow out in the alveoli which can lead to the onset of a life-threatening invasive infection.

In the lung, *A. fumigatus* can invade pulmonary epithelial cells by a process designated as induced phagocytosis (DeHart et al., 1997; Liu et al., 2016; Wasylnka and Moore, 2003). Conidia can survive in the phagosomes of immune cells but also epithelial cells for some time (Amin et al., 2014; Jahn et al., 2002; Schmidt et al., 2020; Seidel et al., 2020; Thywißen et al., 2011). Similar strategies to evade the host’s immune system, (*e.g.* the manipulation of the formation of a functional phagosome), have been described for several bacterial and fungal pathogens (Erwig and Gow, 2016; Flannagan et al., 2012; Schmidt et al., 2020). In many microbial pathogens, this process is linked to the secretion of effector proteins that interfere with the host’s membrane trafficking system (Ledvina et al., 2018; Walpole et al., 2020). For *A. fumigatus*, we previously established that the conidial pigment dihydroxynaphthalene (DHN) melanin prevents the formation of functional phagolysosomes containing conidia (Jahn et al., 2002; Thywißen et al., 2011). Mechanistically, DHN-melanin was found to sequester Ca^2+^ and thus prevents LC3-associated phagocytosis *via* interference with calcium/calmodulin dependent signaling pathways (Kyrmizi et al., 2018). In addition, DHN-melanin reduces the formation of lipid-raft microdomains in the phagolysosomal membrane, which are essential for the generation of a fully functional phagolysosome (Schmidt et al., 2020). Even though the importance of DHN-melanin for this process was established, we were puzzled by the observation that some non-melanized mutant conidia (Δ*pksP*) still escaped killing by phagocytes (Akoumianaki et al., 2016; Schmidt et al., 2020), indicating that additional mechanisms might be involved in the immune escape of this pathogen.

By investigating this phenomenon, we discovered an unprecedented strategy by a fungal pathogen to evade the host immune system by re-directing phagosomes containing conidia to exocytosis. This is based on the specific interaction of the fungal surface protein HscA with the human p11 protein (also called S100A10 or the light chain of annexin A2). As shown here, p11 is a decisive regulatory node for directing endosomes to different pathways.

## Results

### Surface-exposed HscA protein of *A. fumigatus* binds to host epithelial cells

Surface proteins of microbial pathogens play an important role in the host-pathogen interaction, since they are accessible to cellular host proteins. A strategy to identify microbial surface proteins binding to host cells is their specific labeling by biotinylation (Jia et al., 2020) coupled with affinity purification (Liu et al., 2016). To identify such proteins of *A. fumigatus* binding to epithelial cells, we incubated A549 cells with protein extracts containing biotinylated proteins of the fungal surface (Figure 1A). As observed by immunofluorescence imaging of biotin using Alexa Fluor^TM^ 488 streptavidin, we observed that in particular the use of protein extracts of germlings (Gm) led to labeling of the surface of A549 epithelial cells (Figure 1B). This finding suggests that fungal protein(s) binding to A549 cells is (are) relatively abundant on the surface of germlings but not dormant conidia (Dc) or mycelia (Mc). To find out which protein has bound to host cells, we isolated protein extracts of A549 cells after their incubation with *A. fumigatus* protein extracts and performed western blot analysis by using an anti-biotin antibody. This way, we identified a candidate protein with an apparent molecular mass of 70 kDa after co-incubation of A549 cells with protein extract of germlings and a weaker signal for this protein with swollen conidia (Sc) (Figures 1C and 1D). In our previous proteome analysis of surface proteins of *A. fumigatus* during germination (Jia et al., 2020), six proteins with molecular masses between 65 and 75 kDa were detected in extracts of germinating conidia, including four 70 kDa heat-shock proteins. Multiple peptides covering these heat-shock proteins HscA (Afu8g03930/AFUB_083640), Hsp70 (Afu1g07440/AFUB_007770), Ssc70 (Afu2g09960/AFUB_025800), and TktA (Afu1g13500/AFUB_012990) were detected, but only HscA was predominantly identified in samples of germlings (Jia et al., 2020). In agreement, in contrast for example to the *hsp70* gene, the mRNA steady-state level of the *hscA* gene was upregulated in swollen conidia (Figure 1E). The protein level of HscA was also increased in swollen conidia and germlings (Figure 1F). To verify the conidial surface localization of HscA, we generated an *hscA-myc* strain of *A. fumigatus* expressing a Myc-tagged HscA (Figure S1A, S1C, and S1D). As expected, HscA-Myc could be clearly monitored on the surface of germlings by an anti-Myc antibody (Figure 1G). These results suggested that the 67 kDa heat-shock protein HscA from *A. fumigatus* represents a host cell-binding protein.

**Figure 1.**
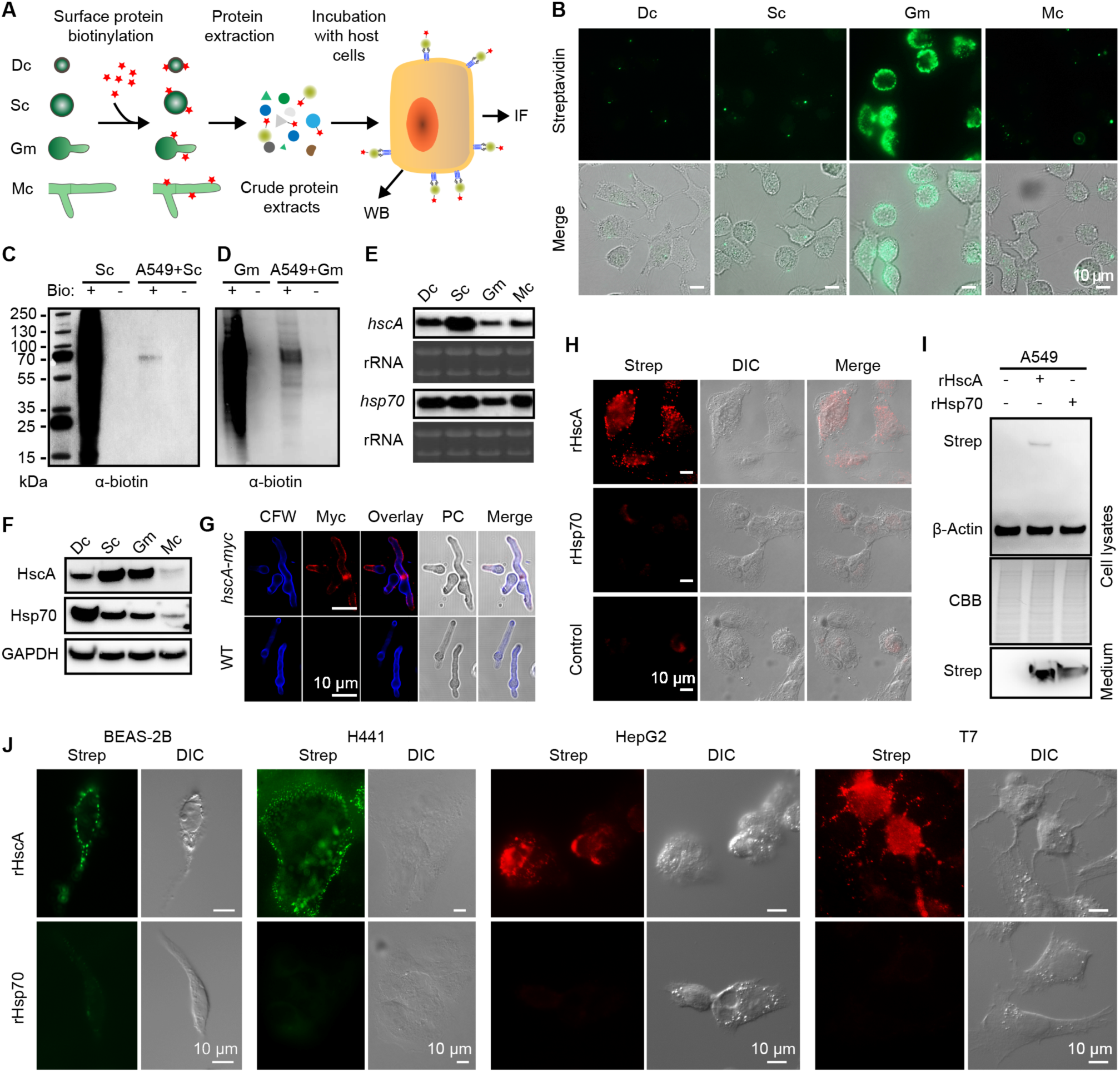
Fungal surface protein HscA binds to the surface of host cells. (A) Scheme illustrating the experimental set-up used for experiments shown in (B–D) Abbreviations: Dc, dormant conidia; Sc, swollen conidia; Gm, germlings; Mc, mycelia; IF, Immune fluorescence analysis; WB, Western blot analysis. (B–D) Biotinylated *A. fumigatus* surface protein binds to the surface of host cells. Surface proteins of Dc, Sc, Gm, and Mc were biotinylated with Ez-Link sulfo-NHS-LC biotin. Subsequently, protein extracts of these different fungal morphotypes were incubated with A549 cells. (B) Proteins bound to cells were detected with Alexa Fluor^®^ 488 Streptavidin by fluorescence microscopy. (C and D) Immunoblot analyses of A549 cells incubated with protein extracts of *A. fumigatus* (C) Sc and (D) Gm by using an anti-biotin antibody. Molecular masses of standard proteins are indicated in kDa on the left margin of (C). Bio +, addition of biotinylated surface proteins. (E) *hscA* expression is up-regulated in Sc. Total RNA of WT *A. fumigatus* Dc, Sc, Gm, and Mc was analyzed using Northern blotting with probes complementary to *hscA* or *hsp70*. rRNA bands are shown as loading control. (F) The HscA protein level is up-regulated in Sc and Gc. Immunoblot analysis of HscA and Hsp70 of WT *A. fumigatus* Dc, Sc, Gm, and Mc using a rabbit anti-HscA antibody and mouse anti-Hsp70 antibody. GAPDH bands are shown as loading control. (G) Detection of HscA on the surface of *A. fumigatus* germlings. Germlings of strain *hscA-myc* and WT were stained with calcofluor white (CFW) and anti-Myc antibody. PC, phase contrast. See also Figures S1C and S1D. (H–J) Recombinant HscA (rHscA) binds to host cells. See also Figures S1G–J. (H) Binding of rHscA to A549 cells was detected by immunostaining of cells with anti- strep antibody. DIC, differential interference contrast. (I) Detection of rHscA from protein extracts of A549 cells. A549 cells were incubated with recombinant rHscA or rHsp70 for 2 hours at 37°C. Recombinant proteins were detected with an anti-strep antibody. CBB, Coomassie brilliant blue staining. (J) Human bronchial epithelial cells (BEAS-2B), human lung epithelial cells (H441), human liver epithelial cells (HepG2), and mouse lung epithelial cells (T7) were incubated with rHscA or rHsp70 followed by staining with an anti-strep antibody All scale bars, 10 μm.

To provide further proof for this conclusion, we produced recombinant HscA (rHscA) and as a control Hsp70 (rHsp70) in *Escherichia coli*. Both recombinant proteins were fused to a Twin-Strep-tag at their N-terminus. By incubation of the purified recombinant proteins with A549 cells, only rHscA, but not rHsp70, bound to A549 cells (Figures 1H and 1I). To further substantiate our finding of binding of rHscA to cells, we generated an *hscA-gfp* strain of *A. fumigatus* (Figures S1A, S1B, and S1F– H). Then, we incubated A549 cells with protein extracts of this *hscA-gfp* strain and, as a control, with protein extracts of strain *ccpA-gfp* that encodes a GFP fusion with another previously identified surface protein of conidia (Voltersen et al., 2018). After immunostaining with an anti-GFP antibody, HscA-GFP was detected on the surface of A549 cells but not CcpA-GFP (Figures S1I and S1J). Although the Hsp70 protein was found on the surface of *hscA-gfp* dormant conidia (Figure S1H), binding of Hsp70 to A549 cells was not observed (Figure S1I), which further underlines the specific binding of HscA but not of Hsp70 to the surface of A549 cells. In addition, we also detected binding of HscA to several other types of epithelial cells, including human bronchial epithelial (BEAS-2B) cells, human lung epithelial (H441) cells, human liver epithelial (HepG2) cells, and mouse type-II lung epithelial (T7) cells (Figure 1J). Collectively, these results showed that surface exposed heat-shock protein HscA of *A. fumigatus* binds to host epithelial cells.

### HscA is an adhesin and intracellular effector protein interfering with the maturation of conidia-containing phagosomes

To identify a possible role of HscA for the interaction of *A. fumigatus* with the host, we generated an *hscA* deletion mutant (Δ*hscA*) (Figures S1A, S1B, and S1D). Δ*hscA* produced slightly smaller colonies on agar plates (Figures S1K and S1L), but showed no obvious defect in sporulation (Figure S1M) and germination of conidia (Figure S1N), or an altered susceptibility against various stressors (Figure S1O). The minor growth defect could be restored by complementation of Δ*hscA* with *hscA-myc* or *hscA* gene (Figures S1K–O).

Based on its host cell binding ability and conidial surface localization, we also assumed an effector function of HscA intracellularly in epithelial cells. To test this assumption, we incubated wild-type (WT), Δ*hscA* and *hscA-myc* conidia with A549 cells. As measured by an LDH release assay, conidia of the Δ*hscA* strain caused significantly less damage to host cells than WT conidia, *i.e*., WT and *hscA-myc* strain, after 20 h of incubation (Figure 2A). Addition of proteins alone of either rHscA or rHsp70 did not increase LDH release, whereas preincubation of Δ*hscA* conidia with the rHscA protein but not with rHsp70 increased LDH release from A549 cells to levels seen with WT conidia. Thus, addition of rHscA complemented the Δ*hscA* phenotype (Figure 2A). This finding suggests that HscA is important for cell invasion and cell damage by conidia. Therefore, we tested the hypothesis whether HscA functions as adhesin by microscopic imaging. In line with our assumptions, the association of conidia of the Δ*hscA* strain with A549 epithelial cells was reduced compared to WT conidia (Figure 2B). This reduction was abolished by the addition of rHscA, but not rHsp70, to the medium (Figure 2B).

**Figure 2.**
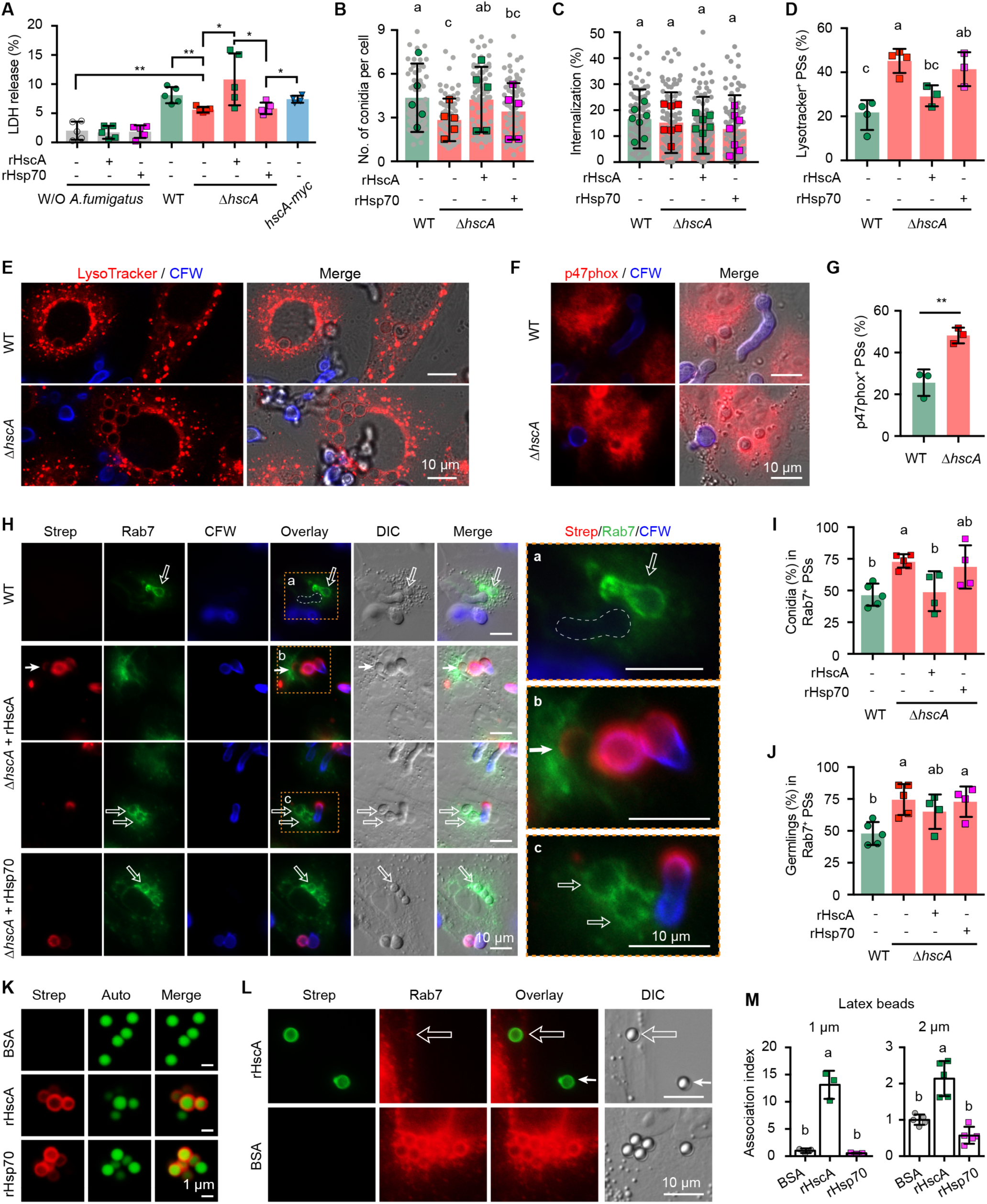
HscA functions as an effector protein. (A) Relative LDH release of A549 cells incubated with conidia of the indicated *A. fumigatus* strain at MOI = 10 for 20 h. Addition of 10 μg/mL rHscA or rHsp70 is indicated. Control cells were incubated with proteins only and without (W/O) *A. fumigatus*. Error bars represent the mean ± SD. \**p* < 0.05, \*\**p* < 0.01 (unpaired, two- tailed t test). (B) Association and (C) internalization of *A. fumigatus* conidia with/by A549 cells. After 8 h of incubation with conidia, A549 cells were washed and extracellular conidia were stained with CFW. Conidia associated with host cells and internalized by host cells were counted. Addition of rHscA and rHsp70 to Δ*hscA* conidia is indicated. Grey dots indicate the calculated values of individual microscopic images of six experiments. Error bars represent the mean ± SD. Colored dots indicate the mean of six individual experiments. (D–G) HscA prevents phagosome maturation. (D) Percentage of acidified phagosomes containing conidia of the WT and the Δ*hscA* strain determined with LysoTracker Red. The addition of proteins is indicated. Microscopy images of (E) LysoTracker^+^ or (F) p47phox^+^ phagosomes of A549 cells containing WT or Δ*hscA* conidia. Extracellular conidia were stained with CFW. (G) Percentage of p47phox^+^ phagosomes of A549 cells containing *A. fumigatus* conidia. Data are mean ± SD. \*\**p* < 0.01 (unpaired, two-tailed t test). (H–J) HscA prevents Rab7 recruitment to phagosomes containing *A. fumigatus* conidia. (H) Immunofluorescence staining of Rab7^+^ phagosomes of A549 cells containing *A. fumigatus* WT or Δ*hscA* conidia. Δ*hscA* conidia were incubated with recombinant HscA (rHscA) or rHsp70 proteins at room temperature for 30 min before inoculation. A549 cells were stained with a mouse anti-Strep-tag antibody and a rabbit anti-Rab7 antibody. Open and thin arrows indicate conidia in Rab7^+^ phagosomes and a Rab7- negative phagosome containing Δ*hscA* conidia coated with rHscA, respectively. Regions labeled with a (WT), b (Δ*hscA* + rHscA), and c (Δ*hscA*) in dashed-line boxes are magnified on the right. Dashed-line circle marks a Rab7-negative phagosome containing germinated WT conidia. (I) Percentage of Rab7^+^ phagosomes of A549 cells containing *A. fumigatus* conidia. (J) Percentage of Rab7^+^ phagosomes of A549 cells containing *A. fumigatus* germlings. (K–M) rHscA prevents Rab7 recruitment to phagosomes that contain latex beads and contributes to association of latex beads with host cells. (K) Immunofluorescence staining of green autofluorescent (Auto) latex beads coated with recombinant proteins rHscA and rHsp70 stained with anti-Strep antibody. Bovine serum albumin (BSA)-coated beads served as negative control. Scale bars, 1 μm. (L) Immunofluorescence staining of A549 cells with phagosomes containing latex beads coated with rHscA or BSA. The open arrow indicates a phagocytosed latex bead coated with rHscA; the small, solid arrow marks an extracellular latex bead. (M) Association index of latex beads with a diameter of 1 μm or 2 μm associated with A549 cells. A549 cells were incubated with latex beads coated with BSA, rHscA or rHsp70, for 8 h at MOI = 20. (B, C, D, G, I, J, and M) Data are mean ± SD; different letters indicate significant difference based on multiple comparisons (Turkey method) after ANOVA. Scale bars represent 10 μm in E, F, H, and L. Abbreviations: LDH, lactate dehydrogenase; PSs, phagosomes; W/O, without; WT, wild type; DIC, differential interference contrast.

As previously shown, *A. fumigatus* conidia are also internalized by alveolar basal epithelial cells (Amin et al., 2014; Keizer et al., 2020; Seidel et al., 2020; Wasylnka and Moore, 2002), and targeted in these cells to phagolysosomes (Amin et al., 2014; Seidel et al., 2020; Wasylnka and Moore, 2003). Therefore, we determined internalization and intracellular processing of conidia in these types of cells. After 8 hours of incubation of conidia with A549 cells, a similar proportion between 12–15% of WT and Δ*hscA* conidia were internalized by A549 cells irrespective of whether rHscA or rHsp70 protein were added to the Δ*hscA* strain (Figure 2C). Interestingly, as indicated by LysoTracker staining, compared to WT conidia about two-fold more Δ*hscA* conidia ended up in acidified phagolysosomes (Figures 2D and 2E). Addition of rHscA, but not rHsp70, reduced the percentage of Δ*hscA* conidia in acidified phagosomes (Figure 2D). Thus, HscA is also involved in inhibiting phagosomal maturation. In addition to acidification, another marker for the maturation of phagosomes is the assembly of the NADPH oxidase complex, consisting of p47phox and other cytosolic subunits, on the phagosomal membrane (Akoumianaki et al., 2016; Kyrmizi et al., 2018; Schmidt et al., 2020). We found that compared to 25% of WT conidia, 48% of Δ*hscA* conidia were localized in p47phox-positive (p47phox^+^) phagosomes (Figures 2F and 2G), which further indicates the importance of HscA for inhibiting the formation of a mature phagolysosome.

To find out by which mechanism HscA modulates phagosome maturation, we analyzed intracellular processing of conidia by A549 cells using immunofluorescence microscopy. We started by analyzing phagosomes for the presence of Rab7 (Figure 2H), which plays an essential role in phagosome maturation (Bucci et al., 2000; Rink et al., 2005; Vieira et al., 2003). While 46% of phagosomes containing WT conidia were Rab7-positive (Rab7^+^), the proportion increased to 73% when phagosomes contained Δ*hscA* conidia (Figure 2I). Addition of rHscA, but not rHsp70, to Δ*hscA* conidia reduced the percentage of Δ*hscA* conidia in Rab7^+^ phagosomes (Figure 2I) most likely due to binding of rHscA to the conidial surface of Δ*hscA* conidia that we had also detected (Figure 2H, b). We also noticed differences in the number of conidia germinating inside phagosomes. Whereas 48% of internalized WT germlings were located in Rab7^+^ phagosomes of A549 cells, this increased to 75% for germinated Δ*hscA* conidia (Figures 2H and 2J). The addition of rHscA or rHsp70 protein did not alter this percentage (Figure 2J), likely because coating with both proteins was restricted to conidia and was also diluted after germination of conidia. Thus, it is apparently a strategy of the fungus to prevent recruitment of Rab7 to phagosomes with HscA on the surface of conidia. Collectively, more WT conidia reside in Rab7- negative phagosomes.

To provide firm evidence that the inhibition of phagosome maturation is due to HscA, we generated latex beads coated with rHscA, rHsp70 or bovine serum albumin (BSA) as control (Figure 2K). A549 cells were incubated with the different latex beads and stained for Rab7. A strong signal of Rab7 was detected on phagosomes containing BSA control beads (Figure 2L), whereas only faint staining of Rab7 on phagosomes containing rHscA beads was observed (Figure 2L). rHsp70 beads were rarely observed attached to or in phagosomes of A549 cells (Figure 2M). In agreement with this finding, coating of Δ*hscA* conidia with rHscA protein blocked recruitment of Rab7 to phagosomes indicated by the lack of staining for Rab7 (Figure 2H). Because internalization of rHsp70 beads was a rare event, we also compared the association of coated 1 µm and 2 µm latex beads with host cells. As expected, rHscA significantly increased the association of latex beads to A549 cells in comparison to rHsp70- or BSA-coated beads (Figures 2M). Overall, these data indicate that HscA mediates adhesion of conidia to host cells and prevents maturation of conidia-containing phagosomes.

### HscA targets the human host p11 protein

To identify a host target protein of HscA, we applied competition binding assays with a human cell surface marker screening kit (BioLegend) combined with imaging flow cytometer (ImageStream X) analysis. Although by using this method no HscA binding receptor was found, we learned that binding of HscA to A549 cells was sensitive to trypsin degradation (Figure S2A) and to fixation of cells with formaldehyde (Figure S2B). These results suggested that HscA binds a proteinaceous partner. To identify such a binding partner of HscA, we applied affinity purification-mass spectrometry. For this purpose, protein extracts of A549 cells were loaded on an rHscA- or rHsp70- loaded Strep-Tactin^®^ (a streptavidin variant) column. Co-purified proteins were analyzed using LC-MS/MS (Figure S2C). About 95 to 226 human proteins were exclusively found in samples co-purified with rHscA or both rHscA and rHsp70 (Figure 3A and Table S1). By comparing the list of proteins, one protein, *i.e*., p11 (also referred to as S100A10), was prominently co-purified (Figure 3A). It was detected in all eluates (n = 5) co-purified with rHscA, three times in the eluates co-purified with rHsp70, and once in the control sample (Figure 3B and Table S1).

**Figure 3.**
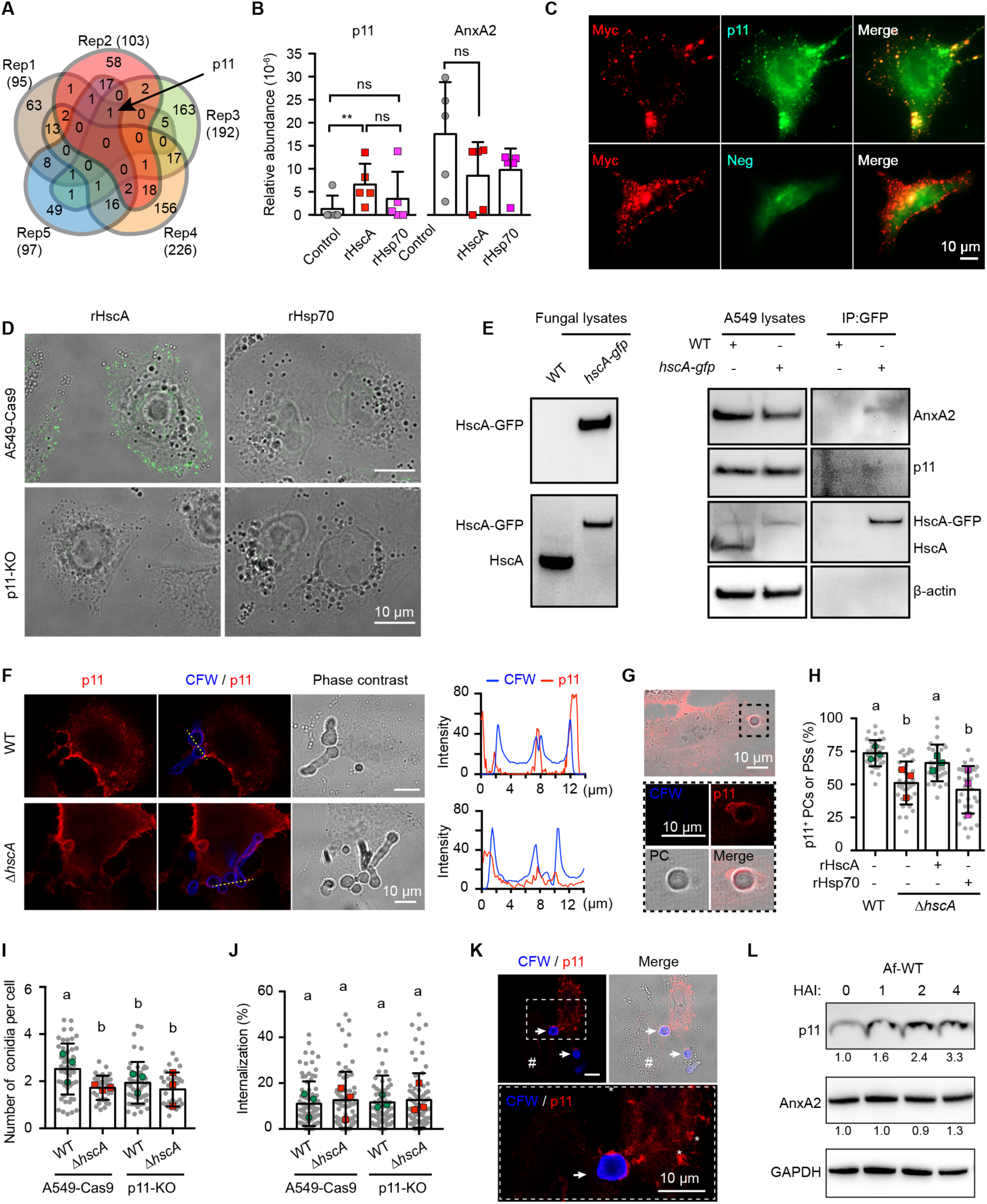
HscA anchors the human p11 protein on phagocytic cups and phagosomal membranes. (A and B) LC-MS/MS detection of p11 co-purified with rHscA. (A) Venn diagram showing the number of proteins (in brackets) detected by LC-MS/MS in eluates with rHscA or in both samples derived from rHscA and rHsp70. Replicates 1–5 are illustrated in different colors and the number of overlapping proteins is marked by numbers. (B) Relative abundance of p11 and AnxA2 co-purified with recombinant proteins rHscA and rHsp70. Data are mean ± SD; **p<0.01; ns, not significant (paired, two-tailed t test). See Figure S2C and Table S1. (C) HscA-Myc colocalizes with p11 on the surface of live A549 cells. A549 cells were incubated with protein extracts from strain *hscA-myc*. Colocalization of HscA-Myc and p11 was indirectly detected using a rabbit anti-Myc antibody and a mouse anti-p11 antibody. For the negative control, the mouse anti-p11 antibody was not added. (D) Binding of HscA to A549 cells is p11-dependent. After incubation with rHscA or rHsp70 for one hour at room temperature, A549-Cas9 or p11-KO cells were stained with mouse anti-Strep antibody. (E) Western blot analysis of p11 and AnxA2 co-purified with HscA-GFP. After co-incubation of A549 cells with protein extracts from *A. fumigatus* WT or *hscA-gfp* strain for 2h at 37°C, cells were washed with PBS and then lysed in IP buffer. GFP-Trap magnetic beads were used to purify HscA-GFP and its binding proteins. Co-purified proteins were analyzed with the antibodies indicated at the right margin. (F and G) p11 is recruited to (F) phagocytic cups and (G) phagosomes of A549 cells. Extracellular conidia were stained with CFW and A549 cells with mouse anti-p11 antibody. Relative signal intensities of the respective emission fluorescence along the lines drawn across the phagocytic cups are depicted on the right of (F). See also Figure S3C and S4A. (H–J) HscA-p11 interaction contributes to adhesion of *A. fumigatus* conidia to host cells. (H) Percentage of p11^+^ structures (phagocytic cups or phagosomes) containing WT, Δ*hscA*, or Δ*hscA* conidia with 10 μg/mL rHscA or rHsp70. (I) Number of WT or Δ*hscA* conidia associated with A549-Cas9 or p11-KO cells. (J) Internalization in % of WT or Δ*hscA* conidia by A549-Cas9 or p11-KO cells. For H– J, grey dots represent counting results of individual microscopy images and colored circles represent means of three individual experiments. Error bars represent the mean ± SD. Different letters indicate significant difference based on multiple comparisons (Turkey method) after ANOVA of the pooled results of individual microscopy images. (K and L) The level of p11 protein increased after *A. fumigatus* infection. (K) Immunofluorescence staining of A549 cells infected with WT conidia of *A. fumigatus* with an anti-p11 antibody. Arrows indicate *A. fumigatus* conidia with induced p11^+^ phagocytic cups. “#” marks a cell with low p11 staining intensity. Asterisks indicate p11^+^ granules. See also Figure S4B. (L) Western blot analysis of p11 and AnxA2 of A549 cells infected with WT conidia at indicated time points. Cell lysates were probed with p11, AnxA2, and β-actin antibodies. Relative band intensity is indicated. See also Figure S4D. All scale bars, 10 μm.

Protein p11 is known to form a heterotetramer (A2t) with Annexin A2 (AnxA2) and is thereby protected from degradation (Gerke and Weber, 1984; He et al., 2008).

AnxA2 binds phospholipids, regulates actin nucleation, and plays important roles in organizing membrane microdomains and vesicle trafficking (Morel et al., 2009). In our affinity purification experiments, however, AnxA2 was detected in all samples irrespective of the presence of rHscA (Figure 3B), suggesting that AnxA2 is not a target of HscA, but potentially the p11 protein. The latter conclusion was further supported by immunofluorescence analysis revealing that HscA-Myc colocalized with p11 on the surface of A549 cells (Figure 3C).

To further substantiate that binding of HscA to host cells is p11-dependent, we generated a p11-knockout A549 cell line (p11-KO) by transfecting a Cas9-expressing cell line with guide RNAs targeting the first CDS of p11 (Figure S3A). DNA sequencing of the generated p11-KO cell line in the region of the gRNA binding region in the p11 gene confirmed frame shifts in both p11 alleles (Figure S3A). The successful generation of a p11-KO cell line was confirmed by Western blot and immunofluorescence analysis demonstrating that in p11-KO cells, the p11 protein was not detectable (Figures S3B and S3C). We then incubated p11-KO cells with the rHscA protein. As expected, rHscA was not detected on the surface of p11-KO cells (Figure 3D).

To further proof that HscA targets p11 in A549 cells, we incubated A549 cells with protein extract from *A. fumigatus* WT or a *hscA-gfp* expressing strain. Similar to the experiments with rHscA (Figure 1I), after incubation, HscA and HscA-GFP were detected in the A549 lysates by Western blots (Figure 3E). Most importantly, p11, together with AnxA2 were co-precipitated with HscA-GFP (Figure 3E). Taken together, these results strongly suggest that p11 and most likely as part of A2t is targeted by the fungal heat shock protein HscA.

### p11 participates in adherence and phagocytosis of *A. fumigatus* conidia

Our data suggest that p11 plays a role in HscA-mediated adhesion and phagocytosis. To further underline this finding, we incubated A549 cells with conidia and examined their interaction microscopically. Surprisingly, p11 and AnxA2 were not only detected on the cytoplasmic membrane, as previously reported (Deora et al., 2004; Fang et al., 2012), but also on both the phagocytic cups (Figures 3F, S3C, S4A) and conidia-containing phagosomes (Figures 3G and S4A). Although p11 was also present on the phagocytic cups containing Δ*hscA* conidia, the overall intensity was much lower compared to the phagocytic cups encasing WT conidia (Figure 3F). Quantification revealed that 74% of WT conidia and only 53% of Δ*hscA* conidia were associated with p11-positive (p11^+^) phagocytotic structures (Figure 3H). This reduction was abolished by addition of rHscA, but not rHsp70, to the medium of A549 cells during their co-incubation with Δ*hscA* conidia (Figure 3H).

Since HscA contributes to the attachment of conidia to host cells (Figure 2B), our findings of accumulation of p11 in phagocytic cups suggested a role of p11 for this process and the phagocytosis of conidia. To address this question, we incubated p11- KO cells with conidia. Consistently, more WT conidia than Δ*hscA* conidia were found associated with A549-Cas9 cells (Figure 3I). In agreement with our assumption, there were less WT conidia associated with p11-KO cells than A549-Cas9 cells. When quantified, we found that the association of WT conidia to p11-KO cells was similar to that measured for Δ*hscA* conidia to A549-Cas9 cells or p11-KO cells (Figure 3I). In agreement with the previous result that HscA did not affect the internalization of conidia in A549 cells (Figure 2C), deletion of p11 did not affect internalization of conidia either (Figure 3J). These results indicate that HscA-p11 interaction contributes to the adherence of *A. fumigatus* conidia to host cells.

### p11 gene expression is induced by *A. fumigatus* infection

After incubation of A549 cells with conidia for 8 hours, the fluorescence signal indicative of the presence of p11 in cells having close contact with conidia was stronger than in cells without conidial contact (Figures 3K and S4B). Furthermore, compared to uninfected A549 cells, a stronger intensity of p11 immunofluorescence and more p11^+^ granules were observed in cells inoculated with *A. fumigatus* conidia (Figures 3K, S4B, and S4C). The increased presence of p11 upon contact with conidia was due to increased p11 levels after incubation of A549 cells with WT conidia and Δ*hscA* conidia, as shown by Western blots (Figures 3L and S4D). This finding indicated that HscA itself is not responsible for the induction of p11 production. The protein level of AnxA2 did not change during the incubation (Figures 3L and S4D), and was not affected by the knockout of p11 (Figure S3B) or knockdown of p11 (Figure S4E). Collectively, our data indicate that the protein level of p11, but not AnxA2, increases upon contact of cells with *A. fumigatus* conidia. In addition, HscA plays a role in anchoring p11 to phagocytic cups and phagosomes of the host cell.

### HscA-induced presence of A2t on phagosomes prevents phagosomal maturation

AnxA2 was previously shown to be present on endosomes and to play a role in early-to-late phagosome transition (Emans et al., 1993; Morel et al., 2009). By contrast, Morel and Gruenberg did not detect p11 on endosomes and postulated its dispensability for AnxA2 association to endosomes. In line with the latter, knockdown of p11 did not affect early-to-late endosomal transition (Morel and Gruenberg, 2007). Here, we found that although AnxA2 was detected on both p11^+^ phagocytic cups and conidia-containing phagosomes (Figures S4A), p11 was only observed on very few AnxA2-positive (AnxA2^+^) phagosomes containing WT conidia (Figure 4A). On phagosomes, indicated by white arrows, both proteins p11 and AnxA2 were present. Compared to WT conidia, more Δ*hscA* conidia were found in AnxA2^+^/p11^-^ phagosomes as indicated by hollow arrows (Figures 4A and 4B). This is in agreement with the observation that in p11 knockdown cells, more than 60% of both WT and Δ*hscA* conidia were found in AnxA2^+^/p11^-^ phagosomes (Figure 4B). Based on this data combined with the high percentage of p11^+^ phagocytic cups containing conidia (Figure 3I), we hypothesize that HscA plays a role in stabilizing A2t on phagosomes.

**Figure 4.**
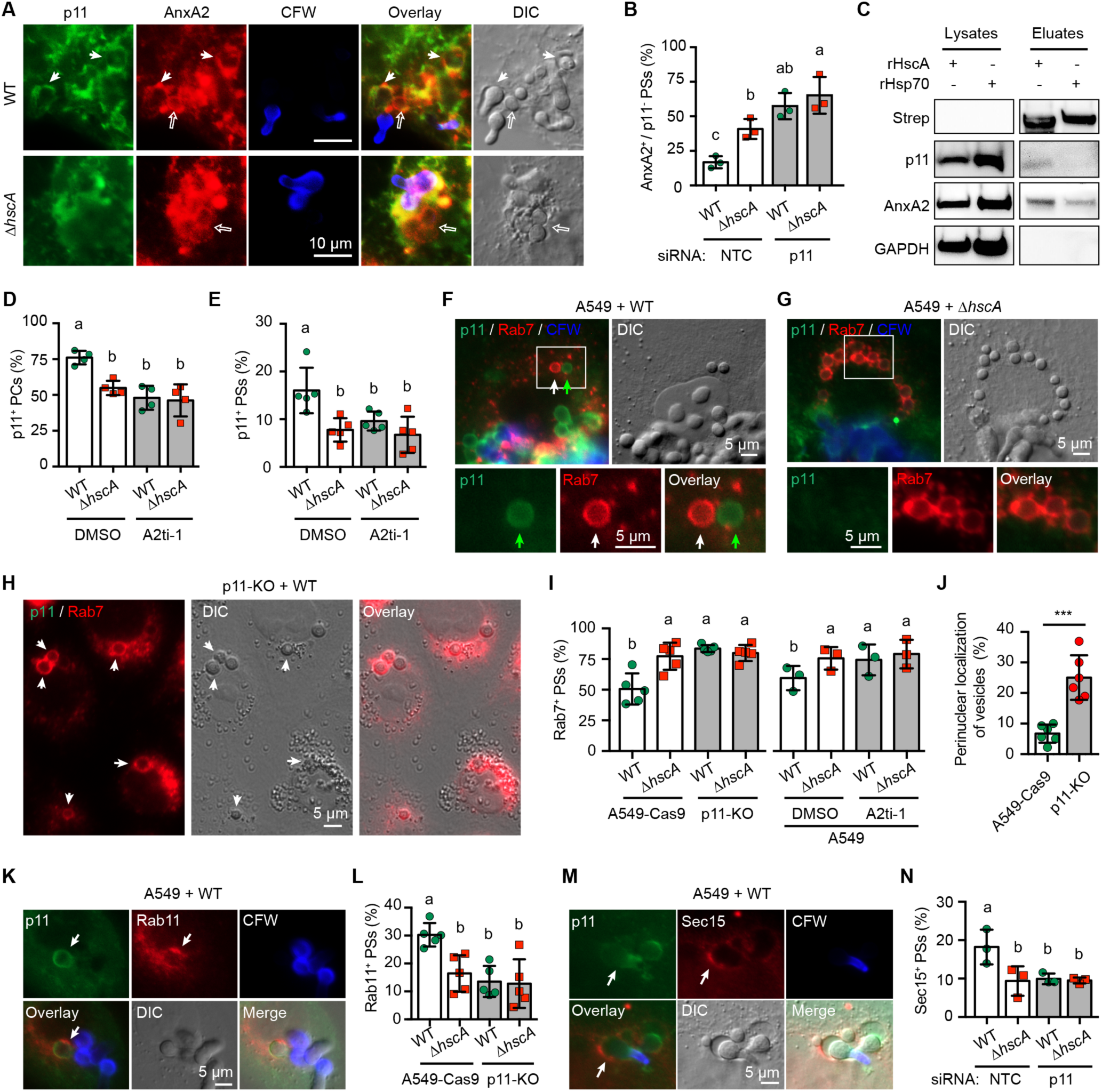
Presence of p11 on phagosomal membrane prevents phagosome maturation. (A–C) p11 colocalizes with AnxA2 on phagosomes containing *A. fumigatus* conidia. (A) Immunofluorescence detection of p11 and AnxA2 of A549 cells after 8 hours of incubation with conidia. A549 cells were washed and stained with CFW, anti-p11, and anti-AnxA2 antibodies. Thin arrows indicate phagosomes that are both p11^+^ and AnxA2^+^, open arrows mark phagosomes that are AnxA2^+^ but p11^-^. Scale bars, 10 μm. See also Figure S4D and S4E. (B) Percentage of AnxA2^+^/p11^-^ phagosomes (PSs) determined by the respective antibodies. A p11 knockdown was generated with p11-targeting siRNA. NTC, non- targeting control RNA. (C) Western blot of lysates of A549 cells and eluates obtained from latex beads coated with the indicated proteins detected by different antibodies. p11 was precipitated with latex beads coated with rHscA. Magnetic latex beads with the size of 1 μm were coated with rHscA or rHsp70 and were incubated with A549 cells for 8 hours. After washing off unbound beads with PBS (pH 7.4), cells were lysed by passing the cells through a 27G needle in homogenization buffer (250 mM sucrose, 3 mM imidazole, pH 7.4). Extracts were probed with the indicated antibodies. (D and E) A2ti-1 inhibits HscA-dependent recruitment of p11 on (D) PCs and (E) PSs. (F–H) Immunofluorescence analyses of A549 cells infected with (F) WT or (G) Δ*hscA* conidia, and (H) p11-KO cells infected with WT conidia. Cells were stained with an anti-p11 (green) and anti-Rab7 (red) antibody, extracellular germlings with CFW. White arrows indicate Rab7^+^ phagosomes containing WT conidia. Green arrows mark a p11^+^ phagosome containing WT conidia. Scale bars, 5 μm. See also Figure S5A. (I) Percentage of Rab7^+^ phagosomes. A549 cells were incubated with 100 µM A2ti-1 or without (only DMSO) for 2 days before incubation with *A. fumigatus* conidia. Furthermore, A549-Cas9 and p11-KO cells were compared. (J) Percentage of accumulated vesicles in the perinuclear region of A549-Cas9 and p11-KO cells. Error bars represent the mean ± SD. ***p<0.001 (unpaired, two-tailed t test). See also Figure S3D. (K–H) Recruitment of Rab11 and Sec15 to p11^+^ phagosomes. (K) Immunofluorescence of p11 and Rab11 in A549 cells after 8 hours of incubation with WT conidia. Arrows indicate phagosomes that are both p11^+^ and Rab11^+^. Scale bar, 5 μm. See also Figure S5B. (L) Percentage of Rab11^+^ phagosomes (PSs). A549-Cas9 cells and p11-KO cells were incubated with WT or Δ*hscA* conidia. (M) Immunofluorescence of p11 and Sec-15 in A549 cells after 8 hours of incubation with WT conidia. Arrows indicate phagosomes that are both p11^+^ and Sec15^+^. Scale bar, 5 μm. See also Figure S5C. (N) Percentage of Sec15^+^ phagosomes (PSs). A549 cells treated with NTC or p11- targeting siRNA were incubated with conidia of the indicated strains. For B, D, E, I, L, and N, data are mean ± SD; different letters indicate significant differences based on multiple comparisons (Turkey method) according to ANOVA.

To test this hypothesis, we incubated A549 cells with magnetic latex beads coated with rHscA or rHsp70 and isolated beads-containing phagosomes as previously described (Goldmann et al., 2021). Immunoblotting confirmed that AnxA2 was eluted from both rHscA- and rHsp70-coated beads, however, p11 was only eluted from rHscA-coated beads (Figure 4C). We also applied the chemical inhibitor A2ti-1 (2-[4- (2-ethylphenyl)-5-o-tolyloxymethyl-4H-[1,2,4]triazol-3-ylsulfanyl]acetamide), that inhibits binding of AnxA2 to p11 (Reddy et al., 2012; Woodham et al., 2015), to A549 cells before infection with conidia. A549 cells treated with A2ti-1 showed a significantly reduced percentage of p11^+^ phagocytic cups (Figure 4D) and phagosomes (Figure 4E) irrespective of the presence of HscA on conidia. Overall, these results indicate that HscA plays a role in stabilizing A2t on phagosomal membranes.

The presence of HscA on conidia reduced staining for Rab7 (Figure 2H–2J), suggesting that interaction of HscA and A2t on phagosomes inhibits phagosome maturation. To test this hypothesis, we analyzed the presence of both p11 and Rab7 on phagosomes by using immunofluorescence. As expected, there was no Rab7 signal detected on p11^+^ phagosomes (Figure 4F). Although a weak p11 signal was still detectable at the interface between the cytoplasmic membrane and extracellular Δ*hscA* germlings, very few Δ*hscA* conidia were located in p11^+^ phagosomes (Figure 4G). Most of the phagocytosed Δ*hscA* conidia located in maturing Rab7^+^ phagosomes that, in addition, were concentrated in the perinuclear region (Figure 4G), where lysosomes accumulate (Korolchuk et al., 2011). To verify the role of p11 or A2t in recruiting Rab7 to phagosomes, we incubated conidia with p11-KO cells or treated A549 cells with the A2t inhibitor A2ti-1. After staining with antibodies against Rab7, in p11-KO cells most of the WT conidia were located in Rab7^+^ phagosomes at the perinuclear region (Figure 4H). Quantification of phagosomes showed that in p11-KO cells 75% of the WT and Δ*hscA* conidia were located in Rab7^+^ phagosomes (Figure 4I). Consistently, a comparable level of Rab7^+^ phagosomes containing WT or Δ*hscA* conidia was observed in A549 cells when treated with A2ti-1 (Figure 4I). These results indicate that A2t prevents phagosomal maturation on phagosomes.

### HscA directs phagosomes to the recycling endosomal pathway and triggers expulsion of conidia

In p11-KO cells we also found accumulation of phagosomes and putative lamellar bodies in the perinuclear region (Figures 4H, 4J, and S3D). This was not the case in wild-type cells, *i.e*., A549 or A549-Cas9 cells (Figures S2B and S3D). Since p11 has been previously shown to play a role in controlling both the distribution of Rab11- positive (Rab11^+^) recycling endosomes (Zobiack et al., 2003) and exosomes release (Chen et al., 2017), we analysed the cell cultures for expulsion of conidia. As shown in Figure S4B, we clearly observed extracellular conidia that could be stained with an anti-p11 antibody. These observations suggest that *A. fumigatus* manipulates the p11- Rab11-recycling of endosomes to escape phagolysosomal killing.

To investigate whether Rab11, which has been shown before to be characteristic of recycling compartments and secretory vesicles (Guichard et al., 2014; Welz et al., 2014), was recruited to p11^+^ phagosomes, we stained A549 cells infected with conidia with an anti-Rab11 antibody. As shown in Figure 4K, Rab11 is indeed present on p11^+^ phagosomes (Figure 4K). Compared to WT conidia, less Δ*hscA* conidia were localized to Rab11^+^ phagosomes of A549-Cas9 cells (Figures 4L). Most importantly, the percentage of Rab11^+^ phagosomes was similar for p11-KO cells infected with WT or Δ*hscA* conidia (Figures 4L), indicating that p11 is essential for HscA-mediated Rab11 recruitment to phagosomes. To substantiate our findings, we analyzed another marker of recycling endosomes and exocytosing vesicles, *i.e*., Sec15, which is an effector of Rab11 (Zhang et al., 2004). As expected, Sec15 was clearly detected on p11^+^ phagosomes (Figure 4M) and, compared to Δ*hscA* conidia, more WT conidia were localized to Sec15-positive (Sec15^+^) phagosomes of A549 cells (Figure 4N). To determine whether this phenotype is directly caused by the presence of HscA, we also incubated A549 cells with rHscA- or rHsp70-coated latex beads. As shown in Figure S5, rHscA-coated beads were found in p11^+^ phagosomes. On the contrary, rHsp70-coated beads were detected in Rab7^+^ phagosomes (Figure S5A). Rab11 and Sec15 were also detected on p11^+^ phagosomes containing rHscA beads, but not rHsp70 beads (Figures S5B and S5C). In summary, these results indicate that WT conidia were more frequently retained in non-matured phagosomes, and could thus be delivered to the recycling endosomal pathway.

Since Rab11 is a marker of recycling endosomes and secretory vesicles, and the cargo within recycling endosomes is targeted to the cell surface, we hypothesized that such cargo conidia might be expelled and leave the cell. To further test this hypothesis, we designed an experiment to check whether conidia could be exocytosed (Figure 5A). Briefly, we incubated A549 host cells with dormant conidia and allowed their ingestion. Next, we added CFW to the medium that exclusively stains conidia outside host cells, as previously reported (Schmidt et al., 2020). After removing CFW, cells were further incubated. Internalized conidia are protected from CFW staining by host cells and can be detected by fluorescence microscopy (Thywißen et al., 2011). As proof for our assumption that conidia are exocytosed, we detected CFW-negative conidia localized to p11^+^ and Rab11^+^ phagocytic cup structures (Figure 5B), suggesting these conidia had been redirected within cells to the surface. This phenotype is dependent on both HscA and p11. About 2.6% of WT conidia and 0.8% of Δ*hscA* conidia were exocytosed by A549 cells. In line, the percentage for WT conidia reduced to 1.2% and was unchanged for Δ*hscA* conidia (1 %) when incubated with p11-KO cells (Figure 5C). This result implies that the interaction of HscA with A2t contributes to the exocytosis of *A. fumigatus* conidia by host cells.

**Figure 5.**
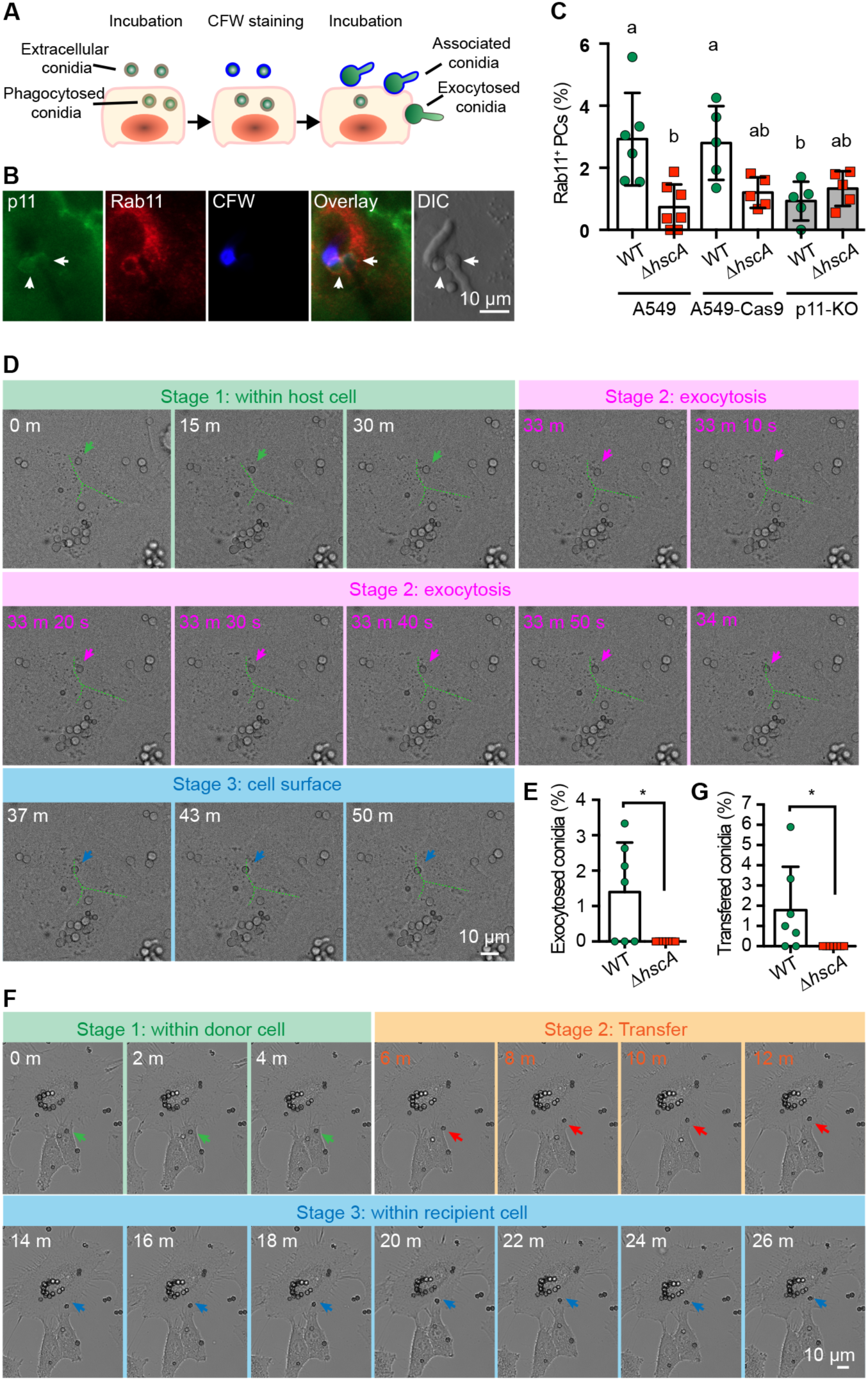
Exocytosis of *A. fumigatus* conidia by host cells. (A) Scheme illustrating the experimental set-up. *A. fumigatus* dormant conidia were incubated with A549 cells for four hours to internalize conidia. Conidia outside of host cells were counter-stained with CFW. After an additional four hours of incubation, cells were fixed, permeabilized and stained with mouse anti-p11 and rabbit anti-Rab11 antibodies. (B and C) *A. fumigatus* conidia escape phagosomes in A549 cells. (B) Immunofluorescence images of potentially exocytosed conidia labeled with anti- p11 (green) and anti-Rab11 (red) antibodies are indicated by arrows. (C) Percentage of *A. fumigatus* conidia attached to host cells with Rab11^+^ PCs. A549 or p11-KO cells were incubated with conidia of indicated strains as described in A. Cells were stained with anti-p11 and anti-Rab11 antibodies. Scale bars, 10 μm. Data are mean ± SD; different letters indicate significant differences based on multiple comparisons (Turkey method) according to ANOVA. (D) Time-lapse image sequence showing exocytosis of WT conidium by A549 cells. The delivered conidium is indicated with arrows with different colors at different stages. Stage 1, the indicated conidium is inside a host cell; stage 2, the conidium is exocytosed to the surface of the host cell; stage 3, the conidium is retained at the host cell surface. Cell borders were indicated with green dotted line. Scale bar, 10 μm. See also Video S1. (E) Percentage of exocytosed conidia of indicated strains from A549 cells. (F) Time-lapse image sequence showing transfer of *A. fumigatus* conidium from donor cell to recipient cell. The delivered conidium is indicated with arrows with different colors at different stages. Stage 1, the indicated conidium is inside a host cell; stage 2, the conidium is transferred to another host cell; stage 3, the conidium is inside the recipient cell. Scale bar, 10 μm. See also Video S2. (G) Percentage of transfer of conidia of the indicated strains from one A549 cell to another cell.

Since the above-mentioned results are deduced from staining of fixed cells, we applied live-cell imaging. As expected, about 1.3% of WT conidia were exocytosed by A549 cells (Figures 5D, 5E and Video S1), while we did not observe such an event for Δ*hscA* conidia (Figure 5E). Interestingly, about 1.5% of WT conidia were transferred from a donor cell to a recipient cell (Figures 5F, 5G and Video S2). Again, this was not observed for Δ*hscA* conidia (Figure 5G). Altogether, these results indicate that targeting of p11 to the conidial surface protein HscA, prevents host phagosome maturation. This allows conidia to germinate or to be redirected to the host cell surface.

### The donor SNP rs1873311 in the p11 gene (*S100A10*) is associated with a decreased risk of invasive pulmonary aspergillosis (IPA) in stem-cell transplant recipients

Our data indicate that p11 plays a major role in phagocytosis and phagosomal processing of conidia. To evaluate this finding in a disease-relevant context, we screened a cohort of hematopoietic stem cell transplant recipients and their corresponding donors for haplotype-tagging single nucleotide polymorphisms (SNPs) in the p11 gene (Figure S6A) and their association with the risk of IPA (Tables S2 and S3). Among the tag SNPs tested, the rs1873311 SNP (T>C), located in the first intron of the p11 gene (Figure 6A), was found to be associated with a reduced risk of IPA. The cumulative incidence of IPA for donor rs1873311 was 25.6 % for T/T and 17.4% for T/C (p = 0.041) genotypes, respectively (Figure 6B and Table S2). The C/C genotype was rare, occurring in only 3 of all 483 donors, and was therefore not plotted. In a multivariate model accounting for patient age and sex, and significant clinical variables (Table S3), the T/C genotype contributed to IPA with an adjusted hazard ratio of 0.86 (95% confidence interval, 0.78–0.96: P = 0.026). We are aware that in humans, in this particular clinical setting, the mechanism proposed appears to be mostly relevant in myeloid cells. Therefore, as a proof of concept, we repeated key experiments in CD45^+^ hematopoietic cells isolated from human lung tissues. Similar to epithelial cells, the isolated hematopoietic cells produce p11^+^ phagosomes and phagocytic cups when confronted with conidia (Figures 6C, 6D, and S7A). There were more WT conidia (22%) than Δ*hscA* conidia (8%) detected in p11^+^ phagosomes (Figures 6E and S7A). Exactly as found in epithelial cells, less WT conidia (43%) than Δ*hscA* conidia (59%) were found in Rab7^+^ phagosomes (Figure 6F). We also incubated the isolated hematopoietic cells with rHscA- or rHsp70-coated latex beads. As shown in Figure S5D, rHscA-coated beads were found in p11^+^ phagosomes, while rHsp70-coated beads were detected in Rab7^+^ phagosomes. These results show that HscA-induced presence of p11 on phagosomes also prevents phagosomal maturation in primary hematopoietic cells.

**Figure 6.**
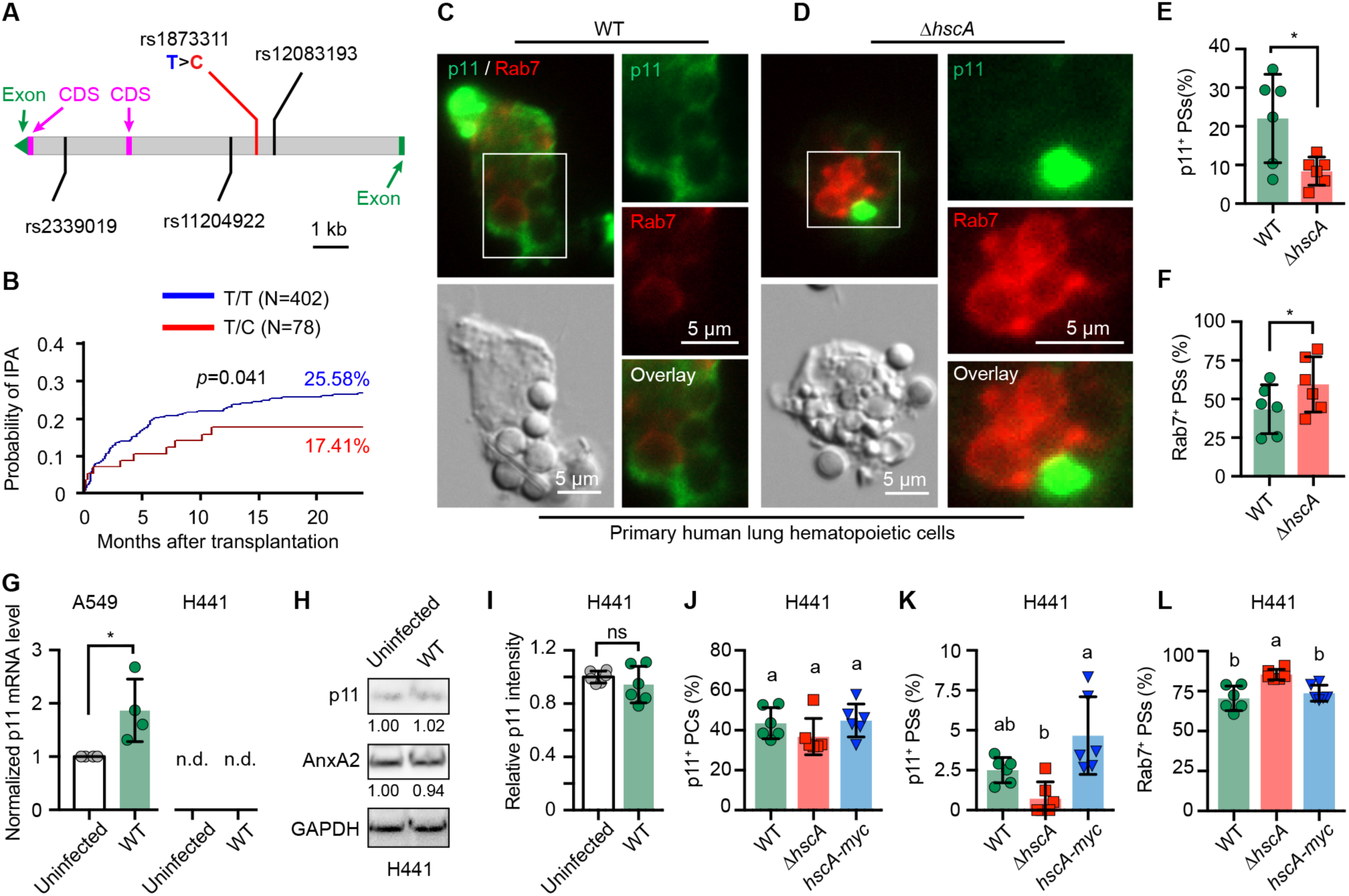
An SNP in the p11 gene (rs1873311) interferes with p11 production and is associated with an increased risk of IPA in stem cell transplant recipients. (A) Position of tag SNPs in human p11 gene. SNP rs1873311 is characterized by a change of thymine (T) to cytosine (C). Introns are indicated with grey blocks, exons are indicated with green blocks, the coding sequences (CDS) of p11 are indicated with magenta blocks. Scale bar, 1 kb. See also Figure S6A and Table S2. (B) Cumulative incidence of IPA in allogeneic stem-cell transplant recipients according to donor rs1873311 genotypes. Data were censored at 24 months. Relapse and death were competing events. *p* value is for Gray’s test. See also Tables S2 and S3. (C–F) p11 excludes recruitment of Rab7 to phagosomes in human lung hematopoietic cells. Hematopoietic cells isolated from human lung tissues were infected with (C) WT or (D) Δ*hscA* conidia for 3 hours. Cells were stained with an anti-p11 (green) and anti- Rab7 (red) antibody. Scale bars, 5 μm. (E and F) Statistical analysis of the percentage of (E) p11^+^ phagosomes (PSs) and (F) Rab7^+^ PSs containing conidia in human lung hematopoietic cells. Data are mean ± SD; *p<0.05 (paired, two-tailed t test). See also Figure S5D. (G) *A. fumigatus* infection increases p11 mRNA level in A549 cells but not H441 cells. qPCR analysis of p11 mRNA level in A549 cells (T/T) and H441 cells (T/C) after co- incubation of WT conidia with cells for 4 hours. Data are mean ± SD; *p<0.05 (unpaired, two-tailed t test). p11 mRNA in H441 cells was not detected (n.d.). See also Figure S7B. (H) p11 protein level in H441 cells is not up-regulated by *A. fumigatus* infection. Western blot analysis of lysates of H441 cells infected with conidia for 4 hours. Extracts were probed with the indicated antibodies. (I) Relative immunofluorescence intensity of p11 induced by conidia in H441 cells for 8 hours. Data are mean ± SD. ns, not significant (unpaired, two-tailed t test). See also Figure S7C. (J–L) Percentage of (J) p11^+^ phagocytic cups (PCs), (K) p11^+^ PSs, and (L) Rab7^+^ PSs- containing conidia in H441 cells after 8 hours of coincubation of cells with conidia of the indicates strains. Data are mean ± SD; different letters indicate significant differences based on multiple comparisons (Turkey method) according to ANOVA. See also Figure S7C.

Since the rs1873311 SNP is not located in the coding region, we hypothesized that it might affect the expression of p11. To address this hypothesis, we tested available human cell lines for the presence of this SNP. DNA sequence analyses at the rs1873311 locus showed that the human cell line A549 displays a wild-type TT homozygous genotype, whereas the human H441 cell line is TC heterozygous (Figure S6B). Thus, Cell line H441 allowed for analyzing the effect of the heterozygous SNP on p11 expression. As shown above, *A. fumigatus* induced up-regulation of p11 both at the mRNA level (Figure 6G) and protein level (Figure 3L) in A549 cells. By contrast, the p11 mRNA was not detectable in H441 cells without or with *A. fumigatus* infection (Figures 6G and S7B). On the protein level, in the H441 cell line the p11 protein was still detectable both by Western blotting and immunofluorescence staining, but its level was not up-regulated by an infection of the cell with conidia (Figures 6H, 6I, and S7C). Overall, these results indicate that the rs1873311 SNP affects the expression of p11 at the transcriptional level and its inducibility by *A. fumigatus*.

This loss of inducibility of the production of p11 protein should be reflected in distinct phenotypes. Therefore, we examined the p11^+^ phagocytic cups induced by conidia and p11^+^ phagosomes in H441 cells because of their heterozygous TC genotype (Figure S7). As shown in Figures 6J and 6K, in H441 cells similar percentages of p11^+^ phagocytic cups (about 40%, Figure 6J) or p11^+^ phagosomes (less than 3%, Figure 6K) were observed with both WT and Δ*hscA* conidia. However, when we compared the percentage calculated for A549 with that for H441 cells, in the heterozygous T/C cell line H441 the calculated percentages were lower than those obtained with the TT homozygous A549 cells (Figures 4D and 4E). When we considered Rab7^+^ phagosomes, slightly less WT conidia (71%) than Δ*hscA* conidia (85%) were detected in H441 cells (Figures 6L and S7C). Comparison of the cell lines, i.e., H441 cells and A549 revealed a much higher percentage of Rab7^+^ phagosomes containing WT conidia in H441 cells than in A549 cells (50%) (Figures 2I and 4I). These findings confirm that a wild-type genotype is required for increased p11 mRNA and protein levels induced by *A. fumigatus*.

## Discussion

The decision whether endosomes enter the degradative or recycling pathway is of fundamental importance for killing of ingested pathogens. Here, we report the discovery of the surface protein HscA of the human-pathogenic fungus *A. fumigatus* that acts as a fungal effector protein influencing this decision. Our data suggest that HscA anchors the human A2t protein complex to phagosomes thereby inhibiting phagosome maturation and inducing expulsion of conidia from epithelial cells (Figure 7). In addition, our data reveal that *A. fumigatus* infection also induces increased levels of human p11 and recruitment of A2t to phagocytic cups and phagosomal membranes independent of the presence of HscA. Anchoring and stabilization of A2t on the phagosomal membrane by HscA directs phagosomes to the secretory pathway by excluding Rab7 but recruiting Rab11 and Sec15 to phagosomes. As a consequence, conidia escape phagolysosomal killing by (a) germination inside a Rab7-negative phagosome, (b) their lateral transfer to other host cells, or (c) by their translocation to the surface of host cells or to the extracellular space. When HscA is lacking as in Δ*hscA* conidia, p11 dissociates from A2t on the membranes of phagocytic cups and phagosomes containing the respective conidia. This leads to (d) recruitment of Rab7 to phagosomes and their maturation to functional phagolysosomes (Figure 7). A strong indication for the clinical relevance of our findings is the identification of an SNP in the non-coding region of the p11 gene that is associated with heightened susceptibility to IPA in hematopoietic stem-cell transplant recipients when present in stem-cell donors. As shown by analyzing this SNP in cell lines we provide evidence that the SNP affects production of the p11 protein.

**Figure 7.**
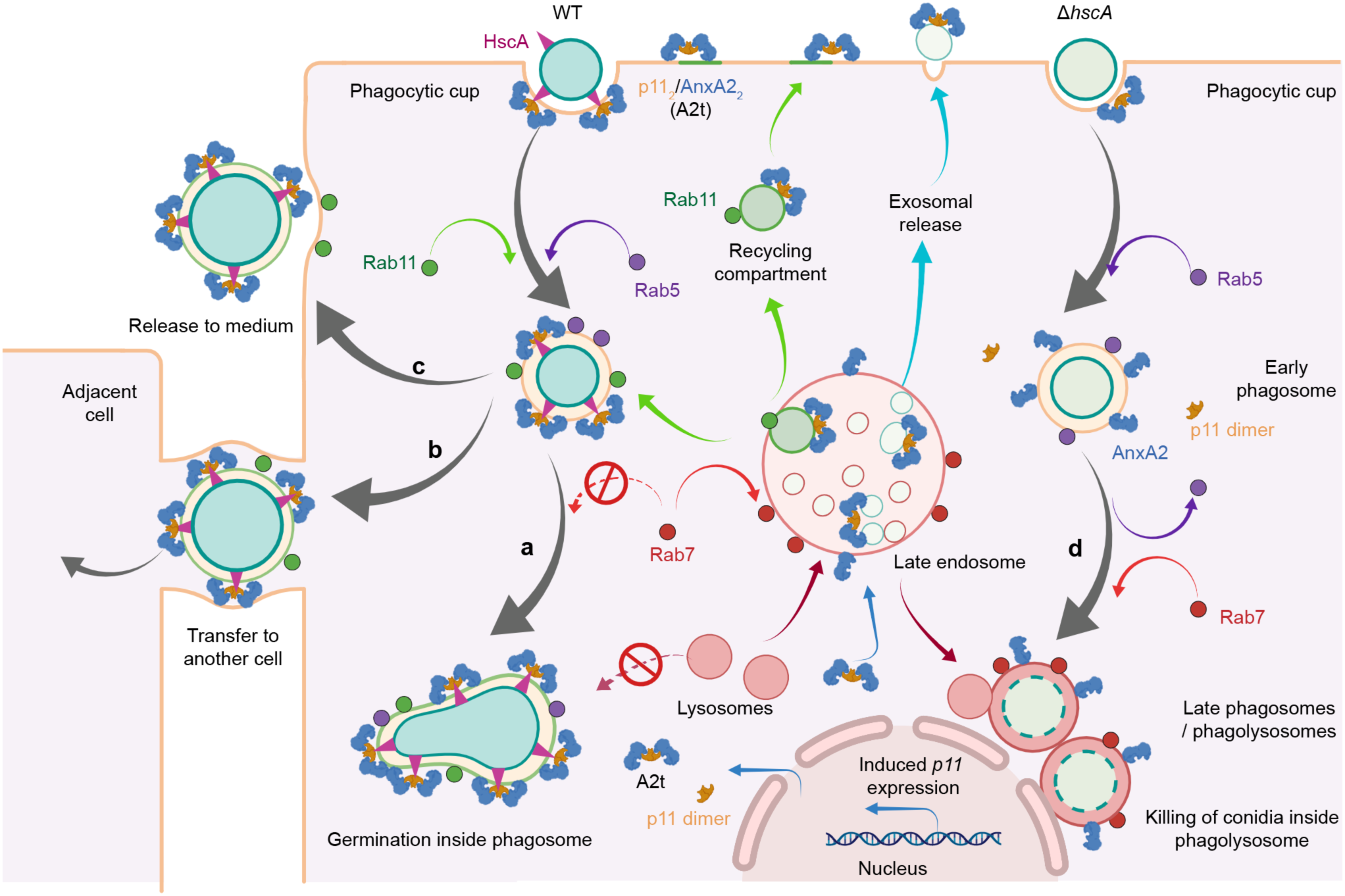
Model of HscA/p11-mediated re-direction of phagosomes to the exocytosis pathway. *A. fumigatus* infection induces the expression of the *S100A10* and the accumulation of its encoded protein p11, which forms heterotetramer (A2t) together with AnxA2, at the phagocytic cups. By binding of conidia to cells, the surface-exposed protein HscA on wild-type (WT) conidia stabilizes the tetrameric complex on the membrane of phagocytic cups and phagosomes. Presence of A2t on phagosomes excludes recruitment of Rab7 to phagosomes and promotes recruitment of Rab11 and Sec15, which are markers of recycling endosomes. As a result, conidia either germinate in the less hostile A2t-positive phagosome (a) or are delivered back to the surface of the host cell. In the latter case, they are either transferred to another cell (b) or released into the medium (c). Without HscA (Δ*hscA*), p11 dissociates from AnxA2 on the phagosomal membranes. Rab7 is recruited to phagosomes and maturation of phagosomes continues to the (d) degradative pathway. Consequently, phagolysosomes concentrate at the perinuclear region where many lysosomes are located (Korolchuk et al., 2011) and conidia are killed. Figure was created with Biorender.com.

Fungal and bacterial pathogens have developed various strategies to escape phagolysosomal killing (Pauwels et al., 2017; Westman et al., 2020). Despite the importance of DHN-melanin on the surface of *A. fumigatus* conidia for delaying phagosome maturation, albino isolates (Couger et al., 2018) and albino Δ*pksP* mutant conidia, that do not contain DHN-Melanin, can still survive macrophage engulfment although with much lower frequency (Kyrmizi et al., 2018). Therefore, it was reasonable to speculate that effector molecules other than DHN-melanin also interfere with phagosomal maturation and thus allow for immune evasion. The here discovered HscA protein represents such an effector protein. HscA was not found in secretome studies of *A. fumigatus* swollen conidia cultivated in RPMI medium (Blango et al., 2020), but traces were detectable in the supernatant of cultures grown with collagen (Shemesh et al., 2017). This finding suggests that the main portion of HscA is bound to the surface of conidia, while a minority may be shedded from the surface.

HscA belongs to the family of HSP70s that are also found on the surface of bacteria, fungi, and mammalian cells (Candela et al., 2010; Sun et al., 2010). In the yeast *Saccharomyces cerevisiae*, putative homologs of HscA are designated as Ssb1 and Ssb2 (HscA has 75% amino acid sequence identity with Ssb2). They are important for folding, co-translational assembly of nascent polypeptides, and also the fidelity of translation termination (Döring et al., 2017; Gribling-Burrer et al., 2019; Willmund et al., 2013). Despite their assumed biological importance, deletion mutants of both *ssb1* and *ssb2* in *S. cerevisiae* (Rakwalska and Rospert, 2004), *hscA* in *Fusarium graminearum* (Liu et al., 2017), *Magnaporthe oryzae* (Yang et al., 2018) and in *A. fumigatus*, as shown here, were viable. Since deletion of *hscA* in *A. fumigatus* only showed a minor phenotype under the conditions tested, the elucidation of the function of the protein other than acting as an effector protein, awaits further studies.

Since Ssb proteins in yeast were shown to bind a large number of substrates (Döring et al., 2017; Willmund et al., 2013), it is reasonable to assume that HscA also binds other host molecules than p11. Among the proteins detected here by affinity purification (Table S1), two microtubule-related proteins DYNLRB2 (Dynein Light Chain Roadblock-Type 2) and MAP4 (Microtubule Associated Protein 4) that are involved in intracellular vesicle trafficking (Pietrantoni et al., 2021; Thapa et al., 2020) were detected. These proteins might also directly or indirectly interact with HscA contributing to further manipulation of endosomal trafficking.

The expression of p11 in various cell types such as BEAS, HeLa, and breast cancer cell line MCF7, was previously shown to be induced by various stimuli, including cytokines, growth factors, dexamethasone or the chemotherapy agent paclitaxel, both at the transcriptional and protein level (Lu et al., 2020; Svenningsson and Greengard, 2007). With *A. fumigatus* conidia, we found another inductor that triggers p11 expression at both transcript and protein level. Because Δ*hscA* conidia still displayed inducing capacity, the p11 induction is independent of HscA. It thus remains to be shown which molecules of *A. fumigatus* trigger expression and in particular translation of p11. For the Δ*hscA* mutant, enrichment of p11 was less prominent, likely because anchoring of p11 by HscA in the phagocytic cup was lacking. As shown here, exocytosis and lateral transfer of conidia are relatively rare events, as similarly observed for the fungal pathogen *Cryptococcus neoformans* and exocytosis for *Candida albicans* (Bain et al., 2012; Ma et al., 2006; Ma et al., 2007). Previously, Shah et al. (2016) observed programmed necrosis-dependent lateral transfer of *A. fumigatus*-containing phagosomes from dying human macrophages to other macrophages. Although β-glucan appears to be a fungal determinant of shuttling, the data suggested that shuttling was driven by a component derived from the conidial cell wall (Shah et al., 2016). At this stage, we cannot exclude that the interaction of HscA with p11 is involved in this process. This, however, seems unlikely because, here, we observed release and transfer of conidia by living cells. Also in zebra fish and mouse phagocytes, a unidirectional shuttling of conidia initially phagocytosed by neutrophils to macrophages was observed that involved phagosome transfer (Pazhakh et al., 2019). This shuttling indeed involved living cells. It remains to be shown whether the same shuttling also holds true for human cells. What appears to be comparable is our observation that conidia are also released and transferred by and to A549 cells when still present in phagosomes, since we detected p11 on the surfaces of exocytosed conidia and germlings.

Until now, molecular mechanism(s) underlying such phenomena on both sides the fungal pathogen and the human host, however, have remained obscured. Here, we found that exocytosis is triggered by the fungal surface molecule HscA that acts as an effector of human p11. The low expulsion rate determined raises the question of the clinical relevance of our findings and a strong indication was the analysis of a cohort of patients at-risk of IPA, which demonstrated a significant association between an SNP at the p11 locus (rs1873311) and heightened susceptibility to infection. In our previous study (Schmidt et al., 2020), the rs3094127 SNP located in the human *FLOT1* gene is lacking in the corresponding gene in mice. Similarly, in this study, there is very low sequence similarity at the rs1873311 SNP locus between the human and mouse p11 gene (Figures S6C–E). Therefore, the SNP cannot be analyzed in mice. Consequently, it is reasonable to assume that there are differences in function between mouse and human p11.

The association with the donor genome appeared to exclude a relevant role of the SNP in the non-hematopoietic compartment, including epithelial cells and endothelial cells although this is a matter of debate. Therefore, we also analyzed hematopoietic cells isolated from human lung tissues and demonstrated identical p11- dependent processing of conidia in these immune cells. In line with the importance of epithelial cell, previously, a comprehensive modelling study revealed the importance of epithelial cells for the defence against *A. fumigatus* infection (Ewald et al., 2021)

The functional relevance of the identified SNP was revealed by comparing the phenotypes of A549 (homozygous T/T) and H441 (heterozygous T/C) cells after infection with *A. fumigatus*. The heterozygous SNP had a major effect on the mRNA steady-state-level of p11, *i.e*., transcript was not detected in H441 cells. After stimulation with *A. fumigatus* conidia, the heterozygous (T/C) SNP at the rs1873311 locus did not lead to upregulated levels of p11 mRNA compared to cells carrying the (T/T) wild-type locus. The identification of this SNP might help to stratify the risk of IPA and identify patients that would benefit the most from antifungal prophylaxis or intensified diagnostics.

A2t and AnxA2 are targeted by various viral, bacterial, and fungal pathogens at different stages of infection (Jolly et al., 2014; Li et al., 2015; Stukes et al., 2016; Taylor et al., 2018b). In the case of the human papillomavirus (HPV), A2t is a central mediator of intracellular trafficking of HPV from early endosomes to late multivesicular endosomes and prevents lysosomal degradation of the virus. Inhibition of A2t by small molecule inhibitor A2ti-1, antibody against p11, or knockout of p11 inhibits HPV infection in host cells (Dziduszko and Ozbun, 2013; Taylor et al., 2018a; Woodham et al., 2015). By using the *anxa2^-/-^*mouse model, AnxA2 has been shown to play important roles against the bacterial pathogen *Pseudomonas aeruginosa* or the fungal pathogen *C. neoformans* (Luo et al., 2016; Stukes et al., 2016). After infection of *anxa2*^-/-^ macrophages with *C. neoformans*, nonlytic exocytosis decreased, whereas the frequency of lytic exocytosis went up and consequently *anxa2^-/-^* mice were more susceptible to *C. neoformans* infection (Stukes et al., 2016). Given that p11 is rapidly degraded in the absence of AnxA2 (Puisieux et al., 1996; Taylor et al., 2018a), a role of p11 or A2t in *C. neoformans* nonlytic exocytosis can be assumed. Here, we found that A2t inhibited not only the maturation of phagosomes containing *A. fumigatus* WT conidia but also contributed to the association of conidia to the surface of host cells. Unlike viruses that need to replicate in the nucleus and avoid entering recycling endosomes (Young et al., 2019), *A. fumigatus* recruits Rab11 to phagosomes in a p11-dependent manner. This prevents phagosomal maturation and, furthermore, triggers the secretory pathway resulting in re-direction of conidia-containing phagosomes back to the cell surface. In this way, fungal conidia germinate and grow in a less hostile environment and can hide from professional phagocytes.

In conclusion, fungal surface protein HscA mediates recruitment and anchoring of A2t to phagosomes. Continuous presence of p11 on the phagosomal membrane puts maturation on hold and redirects phagosomes to the non-degradative pathway. The lack of phagosome maturation allows *A. fumigatus* to grow inside phagosomes and escape cells by outgrowth of germinating conidia and by expulsion and also transfer of conidia between cells. Our finding may contribute to a better understanding of pathogenic mechanisms in many human diseases associated with p11 and AnxA2 such as breast cancer stemness (Lu et al., 2020) and neurological disorders (Jin et al., 2020).

## Acknowledgments

We thank Silke Steinbach, Sylke Fricke, and Flora Rivieccio for excellent technical assistance. We are grateful to Hendrik Huthoff for critical reading of the manuscript. This work was funded by the Deutsche Forschungsgemeinschaft (DFG, German Research Foundation) – the cluster of excellence *Balance of the Microverse* – Project-ID 390713860, Gepris 2051, the DFG-ANR financed French-German project “AfuInf (project number 316898429)”, the DFG Collaborative Research Center (CRC)/Transregio FungiNet 124 ‘Pathogenic fungi and their human host: Networks of Interaction’ (project A1 and Z2; project number 210879364), the DFG CRC 1278 ‘PolyTarget’ (project Z01; project number 316213987), the Leibniz project (K217/2016), the Fundação para a Ciência e Tecnologia (FCT) (PTDC/MED-GEN/28778/2017, PTDC/SAU-SER/29635/2017, UIDB/50026/2020 and UIDP/50026/2020), the Northern Portugal Regional Operational Programme (NORTE 2020), under the Portugal 2020 Partnership Agreement, through the European Regional Development Fund (ERDF) (NORTE-01-0145-FEDER-000039), the European Union’s Horizon 2020 research and innovation programme under grant agreement no. 847507, the “la Caixa” Foundation (ID 100010434) and FCT under the agreement LCF/PR/HR17/52190003, and the Gilead Research Scholars Program – Anti-Fungals.

M.R. and F.S. were members of the excellence graduate school Jena School for Microbial Communication (JSMC) funded by the DFG. C.C. was supported by FCT (CEECIND/04058/2018 to C.C.),

## Author contributions

A.A.B. designed the research project and obtained funding. L.J., O.K., and A.A.B. designed the approach and experiments, and L.J. conducted the majority of the experiments and data analyses. M.R. performed live-cell imaging, confocal microscopy imaging, and qPCR analysis. L.R., M.R., F.S., T.H., and M.S. performed cell culture experiments. P.H. purified recombinant proteins. Z.C. and M.T.F. analyzed the live cell imaging data. C.C., A.C., J.F.L., and A.C.Jr collected patient samples and clinical data, performed the genetic analysis of the patients. T.K. performed LC- MS/MS analysis. B.L. and T.D. collected human lung tissues. L.J., O.K., and A.A.B. wrote the manuscript, and all authors edited the manuscript.

## Data availability

The mass spectrometry proteomics data have been deposited to the ProteomeXchange Consortium *via* the PRIDE (Perez-Riverol et al., 2022) partner repository with the data set identifier PXD030501. All materials within the paper are available from the corresponding author upon reasonable request.

## Methods

### Fungal strains and cultivation

All strains used in this study are listed in Table S4. *A. fumigatus* conidia from WT and knockout strains were collected in water from AMM agar plates after 5 days of growth at 37°C, and were counted using a CASY^®^ Cell Counter, as previously described (Jia et al., 2020). For germination assays, 10^9^ *A. fumigatus* resting conidia were incubated at 37°C in RPMI 1640 (GIBCO) to produce swollen conidia (4 h), germlings (8 h), and hyphae (14 h), as described previously (Jia et al., 2020). For conidia production assay, 10^5^ conidia were spread on AMM agar plates. After incubation at 37°C for 5 days, conidia were collected in 10 mL water and counted using CASY^®^ Cell Counter. For determination of their susceptibility to stressors, serial tenfold dilutions of conidia ranging from 10^5^ to 10^2^ in 1 μL H_2_O were spotted onto AMM agar plates containing 30 μg/mL Congo red, 1 mM 1,4-dithiothreitol, 10 μg/mL tunicamycin, or 0.01% (w/v) SDS. Fungal growth was monitored over time and images were collected before overgrowth of the agar plates. For infection assays, *A. fumigatus* conidia were collected in water from malt agar (Sigma-Aldrich) plates respectively after 7 days of growth at room temperature (22°C). All conidia were harvested in sterile, double-distilled water.

### Strain construction, Southern and Northern blot analysis

A split marker PCR strategy was used to replace the *hscA* gene (AFUB_083640) with the hygromycin B phosphotransferase gene (*hph*) in protoplasts from *A. fumigatus* strain A1160 (CEA17 Δ*akuB*^KU80^) (da Silva Ferreira et al., 2006). Briefly, a 1,085 bp upstream DNA fragment and a 986 bp downstream DNA fragment were amplified from genomic DNA of *A. fumigatus* strain A1160 by high-fidelity PCR using primers HscA- P1, HscA-P2 and HscA-P3, HscA-P4. The two generated DNA fragments were fused with the *hph* cassette (HYG-F and HYG-R) resulting in a 4,809 bp DNA fragment by split marker PCR. A similar strategy was used to generate the *hscA-gfp* strain. A 4,984 bp fragment containing a 1,046 bp 3’ region (without TAA, using primers HscA-P5 and HscA-P6) of *hscA*, a *gfp-ptrA* cassette (using primers PtrA-F, PtrA-R and plasmid pTH1 as template), and a 1,031 bp downstream region (using primers HscA-P7 and HscA-P8) of *hscA* was generated. The *gfp-ptrA* cassette was in-frame fused to *hscA* 3’ region. Then, the fragment was transferred to A1160 protoplasts.

To complement the Δ*hscA* mutant, the intact *hscA* open reading frame, including 1,175 bp of upstream sequence and 683 bp of downstream sequence was amplified from genomic DNA by high-fidelity PCR using primers HscA-Com-F and HscA-Com-R. The resulting DNA fragment was cloned into plasmid pTH1 (Lapp et al., 2014) which was digested with *Kpn*I and *Not*I to obtain plasmid pLJ-HscA-Comp. To generate plasmid pLJ-HscA-Myc, a DNA fragment containing 1,175 bp of upstream sequence and *hscA* without TAA was amplified from genomic DNA by high-fidelity PCR using primers HscA-Com-F and HscA-Myc-R. The DNA fragment was then inserted into plasmid pLJ-Hsp70-Myc (Jia et al., 2020) which was digested with *Kpn*I and *Hin*dIII. Protoplasts of the Δ*hscA* mutant were transformed with plasmids pLJ-HscA-Comp or pLJ-HscA-Myc to generate the respective *A. fumigatus* strains *hscA*c and *hscA-myc*.

For Southern blot analysis, chromosomal DNA of *A. fumigatus* was digested with *Bam*HI. DNA fragments were separated in an agarose gel and blotted onto nylon membranes (Carl ROTH). Northern blot analysis was performed as described previously (Valiante et al., 2016). Total RNA was extracted using a universal RNA purification kit (EURx). 10 μg of RNA was separated on a denaturing agarose gel and transferred onto positively charged nylon membranes (Carl ROTH). Probes were labeled with digoxigenin (DIG) by addition of DIG-11-dUTP (Jena Bioscience) to the PCR mixture. Probe A, synthesized with primers HscA-P8 and oJLJ19-18, probe B, synthesized with primers oJLJ19-45 and oJLJ19-46, were used for Southern blot analysis to verify the *hscA* mutant strain. Probe B was also used for Northern blot analysis to detect *hscA* expression. Probe C was synthesized using primers oJLJ19-33 and oJLJ19-42 to probe *hsp70* mRNA. Probes were detected with an anti-digoxigenin antibody (Roche).

### Cell culture and reagents

Human lung epithelial cells A549 (Cat# 86012804-1VL, Sigma-Aldrich), human distal lung epithelial cells NCI-H441 (Cat# ATCC-CRM-HTB-174D, LGC) were cultured in F- 12K Nut Mix medium (Kaighn’s modification, Gibco) supplemented with 10% (v/v) artificial fetal calf serum (FCS) (HyClone FetalClone III serum, Cytiva). T7 mouse type-II alveolar epithelial cells (Cat# 07021402, ECACC) were cultured in F-12K Nut Mix medium supplemented with 5% (v/v) artificial FCS and with 0.5% (v/v) Insulin-Transferrin-Selenium (Thermo Fisher Scientific). BEAS-2B (Cat# CRL-9609^TM^, ATCC) were cultured in LHC-9 serum free medium (Thermo Fisher Scientific) in flasks precoated with LHC-9 medium supplemented with 0.01 mg/mL bovine fibronectin (Thermo Fisher Scientific), 0.03 mg/mL bovine collagen type I (Sigma-Aldrich) and 0.01 mg/mL bovine serum albumin (BSA; Sigma-Aldrich). HepG2 cells (Cat# ACC 180, DSMZ) were cultured in RPMI-1640 medium supplemented with 10% (v/v) artificial FCS. A549 cells stably expressing Cas9 (Cat# SL504, GeneCopoiea) were cultured as mentioned above for A549 cells, but with addition of 800 µg/mL hygromycin (InvivoGen) as selection marker. All cells were cultivated at 37°C and 5% (v/v) CO_2_.

### Isolation of primary hematopoietic cells from human lung tissues

Healthy human lung tissue was collected during lung surgery on cancer patients (approved by the ethical committee of the Friedrich-Schiller University in Jena, Registration number: 2020-1894 1-Material). Tissue was aseptically removed from the non-tumor affected edges of resected lung wedges or lobes and stored in sterile phosphate-buffered saline (PBS) at 4°C. Tissues were processed between 4 and 24h after surgery. One cm^3^ of the tissue was cut by surgical blade and chopped to smaller pieces by scissors. Enzyme mixture 1 (2 mL TrypLE Trypsin + 0.5 mL Dispase + 3 µL Elastase) or enzyme mixture 2 (2.5 mL Dispase + 3 µL Elastase + 5 µL DNAse) was added in the falcon tube together with tissue and incubated 30 min at 37°C in water bath. After incubation, the mixture was strained through 70 µm and 30 µm cell strainer and washed thoroughly by DMEM/F12 medium (Gibco) and centrifuged at 300 *g* for 10 min at 4°C. Cell pellet was resuspended in 2 mL Red blood cell lysis buffer (Roche) with 5 µL DNAse (1 mg/mL), incubated for 2 min at room temperature and diluted with 6 mL DMEM/F12 medium. After centrifugation at 300 *g* for 5 min at 4°C, cells were resuspended in 500 µL PEB buffer (autoMACS rinsing solution (Miltenyi Biotec) + 5 µL DNase + 0.5% heat-inactivated FCS) with 10 µL FcR blocking reagent (Miltenyi Biotec), 10 µL biotin labeled anti-CD45 antibody (Miltenyi Biotec). After incubation for 30 min at 4°C, cells were washed with 5 mL DMEM/F12 medium and centrifuged at 300 *g* for 10 min at 4°C. Cell pellet was resuspended in 70 µL PEB buffer with 20 µL anti-biotin magnetic beads and incubated at 4°C for 15 min. After another wash with 5 mL media, cells were resuspended in 1 mL PEB buffer and proceeded to magnetic separation on autoMACSpro separator (Miltenyi Biotec) with Possels program. Cells were counted and viability controlled by Trypan Blue staining (1:1 cell suspension + Trypan Blue) and counted on LUNA-FL™ cell counter (Logos Biosystems). Cells (8 × 10^4^) were seeded on fibronectin (bovine, 0.1 mg/mL) coated Millicell EZ _SLIDE_ 8-Well (Merck) in 300 µL media/well (RPMI + 10% FCS + 1% ultraglutamine + 1% Pen/Strep + 1% HEPES + 1% Na-pyruvate + 1% MEM NEAA + 0.1% mercaptoethanol).

### Knockdown and knockout of human p11 gene

ON-TARGETplus Human S100A10 siRNA SMARTpool (Horizon Discovery) was used to knockdown p11 expression. Briefly, 50 µL per well of transfection solution containing 125 nM siRNA and DharmaFECT (1:100) in serum-free F-12 K Nut Mix medium was added to 8-well slides and incubated for 25 min. 4 × 10^4^ cells in 250 µL F-12 K Nut Mix medium supplemented with 10 % (v/v) FCS were then added to the wells and incubated at 37°C with 5% (v/v) CO_2_. The medium was replaced by fresh complete medium on the next day and cells were analyzed on day 3 after transfection by immunoblotting. The A2t inhibitor A2ti-1 (MedChemExpress) was dissolved in DMSO. A549 cells seeded at 3 × 10^4^ cells/well were incubated in an 8-well slide at 37°C with 100 µM of A2ti-1 for 2 days before incubation with *A. fumigatus* conidia. In control experiments, cells were treated with DMSO at the same concentrations used for A2ti-1 delivery.

To generate CRISPR-Cas9 p11 KO cells, A549-Cas9 (GeneCopoiea) cells were transformed with a mixture of sgRNA plasmids, including HCP216549-SG01-3- 10-A, HCP216549-SG01-3-10-B, and HCP216549-SG01-3-10-C (GeneCopoiea, 0.5 µg of each plasmid), using Lipofectamine 3000 reagent (Thermo Fisher Scientific). Colonies derived from single cells were screened for p11 knockout using western blot and immunofluorescence analysis. Knockouts were further confirmed by sequencing the PCR fragment generated using primers oJLJ21-25 and oJLJ21-26. The genotype of A549 cells (T/T, homozygous) and H441 cells (T/C, heterozygous), at the SNP rs1873311 locus was confirmed by sequencing of PCR fragment generated using primers oJLJ21-41 and oJLJ21-42.

### Biotinylation of surface proteins

The surface biotinylation method was applied as described previously (Jia et al., 2020). Briefly, the fungal conidia and mycelium were washed three times with PBS (pH 7.4), and then incubated in 5 ml of PBS containing 5 mg EZ-Link Sulfo-NHS-LC-Biotin (Thermo Fisher Scientific) for 30 min at 4°C. The reaction was terminated by addition of two volumes of 100 mM Tris-HCl (pH 7.4), and the reaction mixture was incubated further for 30 min. Then the samples were washed another three times with PBS. After addition of 1 mL of PBS containing protease inhibitor (Roche) and 500 μL of 0.5-mm-diameter glass beads (Carl ROTH), conidia, germlings, and hyphae were disrupted using a FastPrep homogenizer with the following settings: 6.5 m/s, 3 times for 30 s each time. The samples were then centrifuged at 16,000 × *g* for 10 min at 4°C. Supernatants were collected and their total protein concentration determined by Pierce™ Coomassie Plus™ (Bradford) Protein Assay.

### Production of antibody against HscA

To produce antibody against HscA, two synthesized antigen peptides Cys- TMSLKLKRGNKEKIESALSDA and Cys-DYKKKELALKRLITKAMATR (Figure S1C) were conjugated to KLH carrier and used for raising polyclonal antibody in rabbits (ProteoGenix, France). The detection of HscA using polyclonal antibody was performed by analyzing the protein extracts of *A. fumigatus* WT, Δ*hscA*, *hscA*c, and *hscA-myc* using western blotting.

### Western blotting

For detection of proteins on western blots, whole protein extracts from fungus or host cells were separated on NuPAGE 4%–12% Bis-Tris Gels (Invitrogen) and transferred to 0.2-µm pore size PVDF membranes (Invitrogen) using the iBlot™ 2 Gel Transfer Device (Thermo Fisher Scientific). Membranes were blocked by incubation in 5% (w/v) milk power or 1 × Western Blocking Reagent (Roche) in Tris-buffered saline and 0.1% (v/v) Tween-20 for 1 h at room temperature. Primary antibody incubation was carried out at 4°C overnight. The primary antibodies including the purified rabbit polyclonal anti-HscA antibody (this study, 1:10,000), mouse monoclonal anti-biotin antibody (Thermo Fisher Scientific, 1:2,000 dilution), mouse monoclonal anti-Hsp70 antibody (Thermo Fisher Scientific, 1:1,000 dilution), rabbit polyclonal anti-Myc antibody (Cell Signaling Technology, 1:5,000 dilution), mouse monoclonal anti-Strep antibody (IBA, 1:1,000 dilution), mouse monoclonal anti-GFP antibody (Santa Cruz, 1:1,000 dilution), mouse monoclonal anti-p11 antibody (BD, 1:1,000 dilution), rabbit polyclonal anti-p11 antibody (1:1,000 dilution), rabbit monoclonal anti-AnxA2 antibody (Cell Signaling Technology, 1:2,000 dilution), rabbit monoclonal anti-β-actin antibody (Cell Signaling Technology, 1:2,000 dilution), and mouse monoclonal anti-GAPDH antibody (Proteintech, 1:2,000 dilution) were used. Hybridization of primary antibody with an HRP-linked anti-mouse IgG (Cell Signaling Technology) or HRP-linked anti-rabbit IgG (Abcam) was performed for 1 h at room temperature. Chemiluminescence of HRP substrate (Millipore) was detected with a Fusion FX7 system (Vilber Lourmat, Germany).

### Production of purified recombinant HscA and Hsp70

The coding sequences of *hscA* (with primers AfhscANdeIf and AfhscABamHIr) and *hsp70* (with primers Afhsp70NdeIf and Afhsp70BamHIr) were PCR amplified from *A. fumigatus* cDNA. The generated DNA fragments were cloned into the vector pnEATST (modified pET15b vector encoding an N-terminal Twin-Strep-tag followed by a tobacco etch virus (TEV) protease site). Recombinant proteins were produced in *E. coli* BL21 (DE3) cells (New England Biolabs) by auto induction (Overnight Express Instant TB Medium, Novagen) at 25°C. Bacterial cells were then harvested by centrifugation (10,500 × g) and stored at -80°C. Frozen bacterial cells were resuspended in lysis buffer (100 mM Tris/HCl, 150 mM NaCl, 1 mM AEBSF, 0.5% (v/v) BioLock, pH 8.0) and disrupted at 1000 bar using a high-pressure homogenizer (Emulsiflex C5, Avestin). After centrifugation (48,000 × *g*) and filtration of the lysates through a 1.2 μm membrane, recombinant proteins were purified by affinity chromatography using a 5 mL Strep-Tactin^TM^XT superflow^TM^ high capacity column (IBA). Proteins were eluted from the column with biotin elution buffer (100 mM Tris/HCL, 150 mM NaCl, 1 mM EDTA, 50 mM biotin, pH 8.0). A HiPrep 26/10 Desalting column (Cytiva) was used to transfer fractionated HscA and Hsp70 peaks to storage buffer (20 mM HEPES, 150 mM NaCl, 5 mM MgCl_2_, 10% glycerol (v/v), 1 mM TCEP, pH 7.5).

### Proteomics analysis

To identify the protein(s) from A549 cells binding to HscA, we incubated 10 mg of A549 cell protein extracts with 50 μg purified recombinant HscA protein for 2 h at 4°C. Proteins were then purified using Strep-Tactin^®^XT spin column kit (IBA). Reduction and alkylation of cysteine thiols was performed by addition of 5 mM tris(2- carboxyethyl)phosphine and 6.25 mM 2-chloroacetamide (final concentrations) followed by incubation at 70°C for 30 min. Subsequently, proteins were dried in a vacuum concentrator (Eppendorf) and resolubilized in 50 µL of 100 mM TEAB. Proteolytic digestion was carried out with a trypsin/Lys-C mixture (Promega) incubated for 18 h at 37°C at a protein to protease ratio of 25:1. Tryptic peptides were evaporated in a vacuum concentrator until dryness, resolubilized in 25 µL of 0.05% trifluoroacetic acid in 98:2 H_2_O/acetonitrile (v/v) and filtrated through a 0.2 µm spin filter (Merck Millipore Ultrafree^®^-MC, hydrophilic PTFE) at 14,000 × *g* for 15 min. Filtrated peptides were transferred into HPLC vials and analyzed by LC-MS/MS.

LC-MS/MS analysis was performed on an Ultimate 3000 nano RSLC system connected to a QExactive HF mass spectrometer (both Thermo Fisher Scientific, Waltham, MA, USA). Peptide trapping for 5 min on an Acclaim Pep Map 100 column (2 cm × 75 µm, 3 µm) at 5 µL/min was followed by separation on an analytical Acclaim Pep Map RSLC nano column (50 cm × 75 µm, 2µm). Mobile phase gradient elution of eluent A (0.1% (v/v) formic acid in water) mixed with eluent B (0.1% (v/v) formic acid in 90/10 acetonitrile/water) was performed using the following gradient: 0 min at 4% B, 5 min at 8% B, 20 min at 12% B, 30 min at 18% B, 40 min at 25% B, 50 min at 35% B, 57 min at 50% B, 62-65 min at 96% B, 65.1-90 min at 4% B. Positively charged ions were generated at spray voltage of 2.2 kV using a stainless steel emitter attached to the Nanospray Flex Ion Source (Thermo Fisher Scientific). The quadrupole/orbitrap instrument was operated in Full MS / data-dependent MS2 Top15 mode. Precursor ions were monitored at m/z 300–1500 at a resolution of 60,000 FWHM (full width at half maximum) using a maximum injection time (ITmax) of 100 ms and an AGC (automatic gain control) target of 1 × 10^6^. Precursor ions with a charge state of z = 2– 5 were filtered at an isolation width of *m/z* 2.0 amu for further HCD fragmentation at 30% normalized collision energy (NCE). MS2 ions were scanned at 15,000 FWHM (ITmax = 80 ms, AGC = 2 × 10^5^) using a fixed first mass of *m/z* 120 amu. Dynamic exclusion of precursor ions was set to 20 s. The LC-MS/MS instrument was controlled by Chromeleon 7.2, QExactive HF Tune 2.8 and Xcalibur 4.0 software.

Tandem mass spectra were searched against the UniProt databases (2020/07/13; YYYY/MM/DD) of *Homo sapiens* (https://www.uniprot.org/proteomes/UP000005640) and *Neosartorya fumigata (Aspergillus fumigatus) Af293* (https://www.uniprot.org/proteomes/UP000002530) using Proteome Discoverer (PD) 2.4 (Thermo Fisher Scientific) and the algorithms of Mascot 2.4.1 (Matrix Science, UK), Sequest HT (version of PD2.4), MS Amanda 2.0, and MS Fragger 2.4. Two missed cleavages were allowed for the tryptic digestion. The precursor mass tolerance was set to 10 ppm and the fragment mass tolerance was set to 0.02 Da. Modifications were defined as dynamic Met oxidation, protein N-term acetylation and Met-loss as well as static Cys carbamidomethylation. A strict false discovery rate (FDR) < 1% (peptide and protein level) and a search engine score of >30 (Mascot), > 4 (Sequest HT), >300 (MS Amanda) or >8 (MS Fragger) were required for positive protein hits. The Percolator node of PD2.4 and a reverse decoy database was used for q value validation of spectral matches. Only rank 1 proteins and peptides of the top scored proteins were counted. Label-free protein quantification was based on the Minora algorithm of PD2.4 using the precursor abundance based on intensity and a signal-to-noise ratio>5. Relative abundance of protein was calculated as PSM (peptide spectrum matches)/protein length/total PSM.

### Infection experiments of cells

For infection experiments, *A. fumigatus* conidia were collected in water from malt agar (Sigma-Aldrich) plates respectively after 7 days of growth at room temperature (22°C). A549 and H441 epithelial cells were seeded in Millicell EZ _SLIDE_ 8-Well at a density of 3 × 10^4^ cells per well and incubated overnight at 37°C in a humidified chamber at 5 % (v/v) CO_2_. Conidia were added at a multiplicity of infection (MOI) of 10. Synchronization of infection was achieved by centrifugation for 5 min at 100 × *g*. For immunofluorescence and microscopy, infection of A549 cells and H441 cells was allowed to proceed for 8 h and infection of primary hematopoietic cells was proceeded for 3 h at 37°C in the humidified chamber at 5% (v/v) CO_2_. For LDH release assay, A549 cells were seeded in 24-well plate at a density of 2 × 10^5^ cells per well, and the incubation was extended to 20 h. LDH activity was measured using the CyQuant LDH cytotoxicity assay (Thermo Fisher Scientific) following the manufacturer’s instructions.

### Immunofluorescence and microscopy

A549 cells were seeded in Millicell EZ _SLIDE_ 8-Well at a density of 3 × 10^4^ cells per well 1 day before treatment and processing for immunofluorescence staining. To stain the *A. fumigatus* proteins binding to host cells, living cells were incubated with 20 μg *A. fumigatus* protein extracts or 2 μg purified rHscA or rHsp70 protein at room temperature for 1 h. After three times of washing with PBS, cells were stained with Alexa Fluor 488-conjugated streptavidin (Thermo Fisher Scientific) at room temperature for 1 h to detect biotinylated *A. fumigatus* surface proteins binding to host cells. To stain host cell binding of HscA-GFP protein, cells were incubated with a rabbit anti-GFP primary antibody (Abcam) for 2 h at room temperature and a secondary antibody for 1 h at room temperature in the dark. To stain host cell binding of purified rHscA protein, StrepMAB-Classic DY-488 or StrepMAB-Classic DY-549 (IBA) were used to stain the cells. For staining of phagosomal markers, cells were first incubated with 250 μg/mL calcofluor white for 10 min at room temperature, as described previously (Thywißen et al., 2011), to stain exclusively extracellular conidia. After three washing steps with PBS, cells were fixed for 10 min with 3.7 % (v/v) formaldehyde, membranes were permeabilized for 10 min with 0.1 % (v/v) Triton X-100/PBS and blocked for 30 min with 1 % (w/v) BSA/PBS. Cells were incubated with primary antibodies at 4°C overnight, followed by incubation with secondary goat anti-mouse IgG Alexa Fluor 488 or goat anti-rabbit IgG DyLight 633 (Thermo Fisher Scientific). To determine phagolysosomal acidification, 50 nM LysoTracker Red DND-99 (Thermo Fisher Scientific) was added to A549 epithelial cells 4 h after infection. After another 4 hours of incubation, cells were stained with CFW (Sigma-Aldrich) for 10 min, and fixed for 10 min with 3.7 % (v/v) formaldehyde. Samples were visualized using a Zeiss LSM 780 confocal microscope or a Zeiss Axio Imager M2 microscope and processed with the Zeiss ZEN software.

For quantification, at least 10 individual images of host cells infected with *A. fumigatus* conidia were counted for each experiment of at least three biological replicates. The numbers of extracellular conidia attached to host cells, internalized conidia, phagosomes with positive markers, phagocytic cups, and host cells were counted. The association of conidia with cells was calculated as: (number of internalized conidia + number of conidia attached to the host cells) / number of host cells. The percentage of internalized conidia was calculated as follows: number of internalized conidia / (number of internalized conidia + number of conidia attached to the host cells) × 100. The percentage of phagosomes with a positive marker was calculated as follows: number of phagosomes with a positive marker / number of internalized conidia × 100. The percentage of p11^+^ phagocytic cups was calculated as follows: number of p11^+^ conidia attached to host cells / number of conidia attached to host cells × 100. The percentage of exocytosed conidia was calculated: number of attached conidia with a Rab11^+^ phagocytic cup / number of total conidia × 100.

For live cell imaging, 5 × 10^4^ A549 cells were cultured overnight in eight-well slides (Ibidi) and were infected with *A. fumigatus* conidia at MOI = 5 for four hours.

The cells were then washed with pre-warmed medium and kept inside an incubation chamber at 37°C, 5% (v/v) CO_2_ before carrying out live cell imaging. Confocal time lapse sequences were captured using a Zeiss LSM 780 confocal microscope using a Plan-Apochromat 20x/0.8 M27 objective lens. Images were generated with a 561 nm diode-pumped solid-state laser and collected by the transmitted light photomultiplier tube of the LSM 780 system. Images were collected for 4 hours at 1–10 sec intervals as Z stacks with 2000 nm Z spacing, recording 9–22 confocal slices at each time point. Images consisted of 1024 by 1024 pixels at a voxel size of 415 × 415 × 2000 nm. Quantification of conidia which were exocytosed or transferred between cells were carried out by counting total internalized conidia per replicate.

### Incubation of latex beads with cells

Latex beads were coated as previously described (Dersch and Isberg, 1999). Briefly, 20 μL of bead solution were sequentially washed in 1 mL PBS and 1 mL coupling buffer (0.2 M Na_2_HCO_3_, pH 8.5 and 0.5 M NaCl), and were resuspended in 100 μL of coupling buffer. Purified rHscA and rHsp70 proteins were added in a concentration of 0.5 mg/mL. The suspensions were incubated at 37°C for 30 min. After adding 500 μL of coupling buffer, the suspensions were sonicated for 5 min. For blocking of beads, 500 μL of 10 mg/mL BSA in coupling buffer was added and it was incubated at 37°C for 1 hour. The beads were washed in 1 mL PBS with 10 mg/mL BSA and stored in 200 μL of PBS containing 2 mg/mL BSA at 4°C. Presence of recombinant proteins on the surface of coated beads were verified by detection of Strep tag using immunofluorescence microscopy (Figure 2K).

To quantify the association of latex beads with host cells, fluorescent latex beads (Sigma-Aldrich) coated with recombinant protein were added to A549 cells seeded in Millicell EZ _SLIDE_ 8-Well at a density of 3 × 10^4^ cells per well at MOI = 20. After 8 hours of incubation at 37°C and 5% (v/v) CO_2_, cells were washed three times with PBS and then fixed with 3.7 % (v/v) formaldehyde. The slides were examined using immunofluorescence microscopy. The association index is calculated as number of latex beads per cell divided by the number of BSA-coated latex beads per cell.

To isolate host proteins associated with latex beads, we coated magnetic latex beads (Sigma-Aldrich, 1 μm mean particle size) with rHscA or rHsp70 respectively. The beads were incubated with A549 cells for 8 hours at MOI = 20. After washing-off the unbound beads with PBS, cells were lysed by passing the cells through a 27G needle in homogenization buffer (250 mM sucrose, 3 mM imidazole, pH 7.4), as previously described (Goldmann et al., 2021). After 5 times of washing with PBS, proteins associated with latex beads were eluted in protein loading buffer and analyzed by Western blotting.

### RNA extraction and qPCR analysis

RNA isolation from cells was performed using the Universal RNA purification Kit (Roboklon GmbH, Berlin, Germany). 3 × 10^5^ of A549 or H441 cells were co-incubated with *A. fumigatus* conidia with MOI = 10 for 4 hours at 37°C and 5% (v/v) CO_2_. Cells were lysed in 400 μL buffer RL containing 10% (v/v) β-mercaptoethanol. RNA was extracted following manufactureŕs protocol for cell culture RNA purification. Complementary DNA (cDNA) was synthesized from RNA using the Maxima H Minus First Strand cDNA Synthesis Kit (Thermo Fisher Scientific). Real-time qPCR was performed using iTaq^TM^ Universal SYBR^®^ Green Supermix (Bio-rad) on a QuantStudio3 real-time PCR system (Thermo Fisher Scientific) with the following thermal cycling profile: 95°C for 20 s, followed by 40 cycles of amplification (95°C for 5 s, 58°C for 34s). 18s ribosomal RNA was used as an endogenous control for normalization.

### Co-immunoprecipitation

One 182 cm^2^-flask of 80% confluent A549 cells was incubated with 20 mg crude protein extract of *A. fumigatus* for 2 h at 37°C. After 5 times of washing with PBS, cells were incubated with 1 mM DSP (dithiobis(succinimidyl propionate), Thermo Fisher Scientific) at room temperature for 30 min. The reaction was quenched with 10 mM Tris-HCl (pH 7.4), and the cells were lysed in 500 μL of IP lysis buffer (Thermo Fisher Scientific) with protease inhibitor (Roche). The lysates were centrifuged at 16,000 × *g* for 10 min at 4°C. The HscA-GFP protein was precipitated with GFP-trap magnetic agarose (ChromoTek). The co-immunoprecipitation of p11, AnxA2, and HscA-GFP was analyzed via western blotting.

### Genetic association study

The genetic association study with IPA was performed in a total of 483 hematological patients of European ancestry undergoing allogeneic hematopoietic stem-cell transplantation at Instituto Português de Oncologia, Porto, and at Hospital de Santa Maria, Lisbon was enrolled in the IFIGEN study between 2009 and 2016. The demographic and clinical characteristics of the patients are summarized in Table S3. Cases of probable/proven IPA were identified according to the standard criteria from the European Organization for Research and Treatment of Cancer/Mycology Study Group (EORTC/MSG) (De Pauw et al., 2008). Patients diagnosed with “possible” invasive fungal infection or with a pre-transplant infection were excluded from the study. Approval for the IFIGEN study was obtained from the SECVS (no. 125/014), the Ethics Committee for Health of the Instituto Português de Oncologia - Porto, Portugal (no. 26/015), the Ethics Committee of the Lisbon Academic Medical Center, Portugal (no. 632/014), and the National Commission for the Protection of Data, Portugal (no. 1950/015). Experiments were conducted according to the principles expressed in the Declaration of Helsinki, and participants provided written informed consent.

### SNP selection and genotyping

Genomic DNA was isolated from whole blood using the QIAcube automated system (Qiagen). SNPs were selected based on their ability to tag surrounding variants with a pairwise correlation coefficient r^2^ of at least 0.80 and a minor allele frequency ≥5% using publicly available sequencing data from the Pilot 1 of the 1000 Genomes Project for the CEU population. Genotyping was performed using KASPar assays (LGC Genomics) in an Applied Biosystems 7500 Fast Real-Time PCR system (Thermo Fisher Scientific), according to the manufacturer’s instructions.

### Statistical analysis

Statistical analysis was performed using Prism 7. One-way ANOVA was used to analyze experimental data with more than two experimental groups followed by Tukey’s multiple comparisons test. Two-tailed unpaired Student’s t test was additionally used for data analysis. The probability of IPA according to *S100A10* genotypes was determined using the cumulative incidence method and compared using Gray’s test (Gray, 1988). Cumulative incidences at 24 months were computed with the *cmprsk* package for R version 2.10.1 (Scrucca et al., 2007), with censoring of data at the date of last follow-up visit and relapse and death as competing events. All clinical and genetic variables achieving a p-value ≤0.15 in the univariate analysis were entered one by one in a pairwise model together and kept in the final model if they remained significant (p˂0.05). Multivariate analysis was performed using the subdistribution regression model of Fine and Gray with the *crr* function for R (Scrucca et al., 2010).

## Supplemental information

**Figure S1. Verification and phenotypic analysis of *A. fumigatus hscA* mutant strains (see also** **Figure 1**).

(A) Scheme of the organization at the chromosomal *hscA* locus of the different *A. fumigatus* strains. Size of generated DNA fragments by *Bam*HI restriction and binding sites of hybridization probe A and B are indicated. *ptrA*, pyrithiamine resistance gene

(B) Southern blot analysis of chromosomal DNA cut by *Bam*HI to confirm the generated recombinant *A. fumigatus* strains. A DNA band obtained with probes A and B with the size of 6.3 kbp is characteristic of the WT strain, a band obtained with probe A with the size of 4.7 kbp of Δ*hscA* strain and 5.3 kbp of *hscA-gfp* strain. A band obtained with probe B with the size of 3.1 kbp is indicative of both strain *hscA-gfp* and *hscA*c.

(C) Schematic representation of HscA-Myc substrate binding domains (SBDs) consisting of SBD-β (purple) and SBD-α (lid domain, red) (Gumiero et al., 2016). Myc tag (M) was fused to the C-terminus of SBD. Positions of antigens 1 and 2 used for polyclonal antibody generation are marked with red lines. Lysine residues with biotinylation marks are indicated (Jia et al., 2020).

(D) Western blot of protein extracts from dormant conidia of indicated strains with antibodies against HscA, Myc-tag, Hsp70, or GAPDH. Conidia were harvested from malt agar plates after 7 days of cultivation at 22°C.

(E) Western blot of HscA in dormant conidia. Strains *hscA-myc* and *hscA-gfp* were inoculated on AMM or malt agar and incubated at 22°C or 37°C for 7 days. Protein extracts from dormant conidia were probed with anti-Myc, anti-GFP or anti-GAPDH antibodies. See also Figure 1F and 1G.

(F) Western blot for the detection of the HscA-GFP fusion protein with an anti-GFP antibody.

(G) DNA sequence of the fusion site of genes *hscA* and *gfp* present in the *hscA-gfp* strain. The DNA fragment containing the 3’ region of *hscA* and the 5’ region of *gfp* was PCR amplified using the primer pair oJLJ19-45 and oJLJ18-57, and then sequenced using primer oJLJ19-45.

(H) Immunofluorescence staining of Hsp70 localized on the surface of *hscA-gfp* dormant conidia with the anti-Hsp70 antibody. Conidia incubated with secondary antibody served as negative control.

(I) Immunofluorescence staining of HscA-GFP binding to A549 cells. A549 cells were incubated with protein extracts of strain *hscA-gfp* at room temperature for 1 h. Cells were then incubated with anti-GFP antibody or anti-Hsp70 antibody. A549 cells without incubation with fungal protein extracts served as negative control.

(J) Immunofluorescence staining of HscA-GFP binding to A549 cells. A549 cells were incubated with protein extracts of strains *hscA-gfp* or *ccpA-gfp* at room temperature for 1 h. Cytoplasmic membrane of A549 cells was stained with Oregon Green™ 488 conjugated wheat germ agglutinin (WGA). Protein was stained with anti-GFP antibody. For H–J, goat anti-rabbit IgG Dylight 633 or goat anti-mouse IgG Dylight 633 were used to detect primary antibodies. Scale bars, 10 μm.

(K–O) *hscA* gene deletion caused no severe growth defects.

(A) (K) Images of serial 10-fold dilutions of conidia of the indicated *A. fumigatus* strains inoculated onto AMM agar and incubated for 4 days at 22°C, or 2 days at 37°C or 42°C.

(B) (L) Colony diameter of indicated strains. 10^5^ conidia were inoculated at the center of AMM agar plates and incubated at the indicated temperatures for 3 days.

(C) (M) Number of conidia of indicated strains on AMM agar plates after 3 days of incubation at 37°C. 10^5^ conidia were freshly harvested and spread onto AMM agar plates. Conidia were harvested from each agar plate with 10 mL of sterile water.

(D) (N) Number of germlings of the indicated *A. fumigatus* strains incubated in RPMI medium for 8 hours at 37°C.

Data are mean ± SD; different letters indicate significant differences based on multiple comparisons (Turkey method) according to ANOVA.

(A) (O) Images of serial 10-fold dilutions of conidia of the indicated strains that were spotted on AMM agar plates containing 30 μg/mL Congo red, 1mM DTT, 10 μg/mL tunicamycin, or 0.01% (w/v) SDS at 37°C for 2 or 3 days.

**Figure S2. Identification of potential binding partners of HscA (see also** **Figure 3). (A and B) Pre-treatment of A549 cells with (A) trypsin or (B) formaldehyde abolished binding of HscA to A549 cells.**

(A) Immunofluorescence staining of A549 cells incubated with protein extract of dormant conidia for 1 h at room temperature and stained with anti-GFP antibody after pre-treatment of A549 cells with trypsin. A549 cells were suspended in enzyme-free dissociation buffer or trypsin digestion buffer.

(B) Immunofluorescence of A549 cells pre-fixed with 4% (v/v) formaldehyde in PBS. Then, cells were incubated with rHscA or protein extracts from strains *hscA-gfp* or *ccpA-gfp* for 1h at room temperature followed by detection using indicated antibodies. All scale bars, 10 μm.

(C) SDS-PAGE of A549 protein extracts incubated with the indicated recombinant proteins. A549 cell lysates were incubated in IP buffer, with rHscA or rHsp70 for 2 h at 4°C. Samples were purified by Strep-Tactin^®^XT spin columns, and then analyzed by LC-MS/MS. Molecular masses of standard proteins indicated on the left side.

**Figure S3. Knockout of the p11 gene in A549 cells** (see also Figure 3).

(A) Verification of generated p11-KO cell line by DNA sequencing. A 704 bp DNA fragment was amplified from A549-Cas9 cell line or p11-KO cell line and sequenced using primers oJLJ21-25 and oJLJ21-26. The DNA fragment obtained from p11-KO cells was further cloned into pJET1.2 for DNA sequencing. As indicated with red boxes, deletion of ten base pairs in allele 1 and two single base pairs in allele 2 causes a frame shift of the p11-coding sequence and results in a p11 knockout.

(B) Western blot of protein extracts of A549 cell lines A549-Cas and p11 knockout p11-KO with antibodies against p11, AnxA2, or β-actin.

(C) Immunofluorescence staining of p11 in cell lines infected with WT conidia. Arrows indicate p11^+^ phagocytic cups. See also Figure 3F.

(D) Microscopic image of perinuclear localization of vesicles in p11-KO cells. See also Figure 4H.

Scale bars, 10 μm.

**Figure S4. p11 protein level increases by *A. fumigatus* infection (see also** **Figure 3** **and** **Figure 4**).

(A) Immunofluorescence staining of AnxA2 and p11 on phagocytic cups (upper row) and phagosomes (bottom row) of A549 cells infected with WT conidia for 8 h. Arrows indicate a phagosome containing conidia. Extracellular conidia and germlings were stained with CFW. Cells were stained with anti-p11 and anti-AnxA2 antibodies. See also Figures 3F and 4A.

(B–D) p11 protein level is upregulated after *A. fumigatus* infection.

(A) (B) Immunofluorescence staining of p11 in A549 cells infected without or with *A. fumigatus* for 8 hours at MOI = 10. Arrows indicate an extracellular germling with p11 staining. Extracellular *A. fumigatus* conidia and germlings were stained with CFW. Cells were stained with anti-p11 antibody. See also Figure 3K.

(B) Relative immunofluorescence intensity of p11 induced by fungal infection. Error bars represent the mean ± SEM. *p<0.05 (unpaired, two-tailed t test).

(C) Western blot showing induction of p11 protein expression by Δ*hscA* conidia or IFN- γ. Incubation of A549 cells with Δ*hscA* conidia (MOI = 10) or IFN-γ (50 ng/mL) for indicated time; cell lysates were analyzed by Western blot analysis and probed with anti-p11, anti-AnxA2 or anti-β-actin antibodies. See also Figure 3L.

(D) Knockdown of p11 expression in A549 cells with p11-targeting siRNA. After incubation of cells with indicated concentrations of p11-targeting siRNA or non-targeting control siRNA (NTC, 25 nM) for 48 h, cell lysates were probed with anti-p11, anti-AnxA2 or anti-GAPDH antibodies.

Scale bars, 10 μm.

**Figure S5. Differential recruitment of p11 and Rab proteins to phagosomes** (see also Figure 4 and Figure 6).

(A–C) Recruitment of phagosomal markers to latex beads with a diameter of 3 μm coated with protein rHscA or rHsp70. After incubation of A549 cells with the beads for 8 h, cells were fixed, permeabilized, and stained with the indicated antibodies: (A) anti- p11 and anti-Rab7; (B) anti-p11 and anti-Rab11; (C) anti-p11 and anti-Sec15.

(A) (D) HscA recruits p11 to and excludes Rab7 from phagosomes containing latex beads in macrophages. After incubation of primary lung macrophages with beads for 3 h, cells were fixed, permeabilized, and stained with an anti-p11 antibody and an anti-Rab7 antibody. See also Figure 6C and 6D.

Scale bars, 10 μm. White arrows indicate positive staining of phagosomes by indicated antibodies.

**Figure S6. Sequence analysis of SNPs in *S100A10* (p11) gene (see also** **Figure 6**).

(A) Graphical view of the haplotype-based tagging strategy for the SNPs in the *S100A10* gene. SNPs with a minor allele frequency (MAF) above 0.05 were selected from the publicly available sequencing data from the Pilot 1 of the 1000 Genomes Project for the CEU population (Northern Europeans from Utah). Tag SNPs are indicated with the red circles and nsSNPs are indicated with blue circles. Linkage disequilibrium (LD) values were used to define LD blocks tagged by each SNP. See also Figure 6A.

(B) DNA sequences of cell lines A549 and H441 at the rs1873311 SNP locus. DNA fragments were PCR amplified using primers oJLJ21-41 and oJLJ21-42. The sequence shows the C/T heterozygous genotype of the H441 cell line.

(C–E) Alignment of human and mouse p11 gene.

(A) (C) Signatures of human and mouse p11 gene. Introns are indicated with grey blocks, exons with green blocks, the coding sequence (CDS) of p11 is indicated with magenta blocks. Scale bar, 1 kilo base (kb).

(D and E) Alignment of p11 DNA sequences at the locus (D) rs12083193 and (E) rs1873311. Asterisks indicate aligned bases with same identity. The identity of aligned sequences labeled with the same colored box is indicated as regional sequence identity.

**Figure S7. Immunofluorescence staining of (A) primary human hematopoietic cells and (B) H441 cells infected with *A. fumigatus* (see also** **Figure 6**).

(A) Hematopoietic cells isolated from human lung tissues were infected with WT or Δ*hscA* for 3 h. Cells were stained with anti-p11 antibody. See also Figure 6C and 6D.

(B) p11 mRNA was not detected in H441 cells. cDNA of H441 cells or A549 cells were PCR amplified using primers of the TBP gene (encoding the TATA-box binding protein) as a positive control and primers of the p11 gene. See also Figure 6G.

(C) H441 cells were infected with WT, Δ*hscA,* or *hscA-myc* conidia for 8 h. Extracellular conidia and germlings were stained with CFW. Cells were stained with anti-p11 and anti-Rab7 antibodies. White arrows indicate p11^+^ phagosomes and yellow arrows indicate p11^+^ phagocytic cups containing conidia. Red arrows indicate Rab7^+^ phagosomes. Scale bars, 10 μm. See also Figures 6I–L.

**Table S1. Human proteins identified using affinity purification mass spectrometry.** (.xlsx)

Table S2. Frequency of *p11* (*S100A10*) genotypes among cases of IPA and controls, and association test results. (.docx)

**Table S3. Baseline characteristic of transplant recipients enrolled in the study.**

(.docx)

**Table S4. Key resources used in this study.** (.xlsx)

Video S1. Exocytosis and phagocytosis of *A. fumigatus* conidium by A549 cells. Video S2. Shuttling of *A. fumigatus* conidium from donor cell to recipient cell.

