## Supplementary figures and images for "Re-direction of phagosomes to the recycling expulsion pathway by a fungal pathogen"

### Supplementary Figures S1-S7

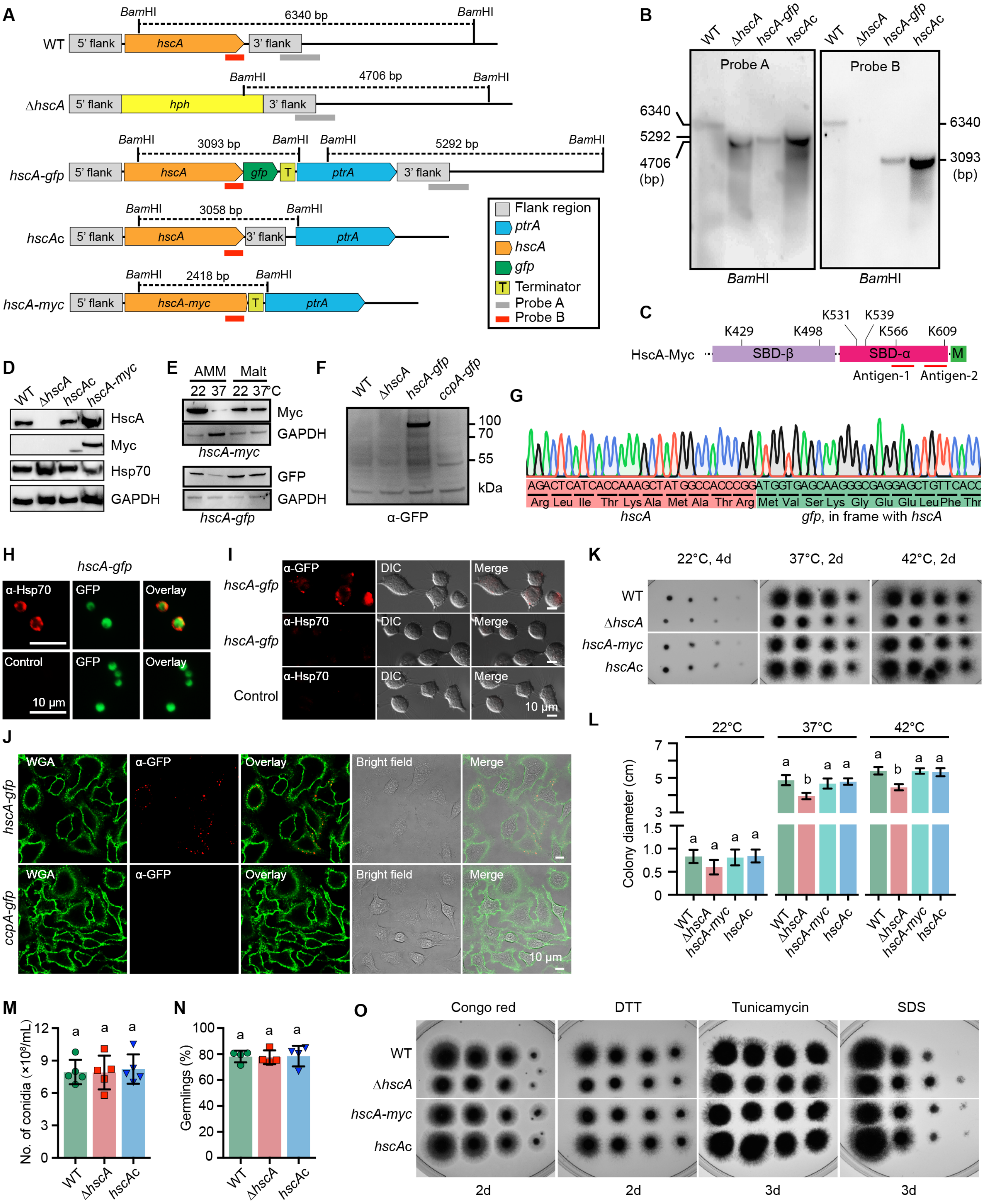

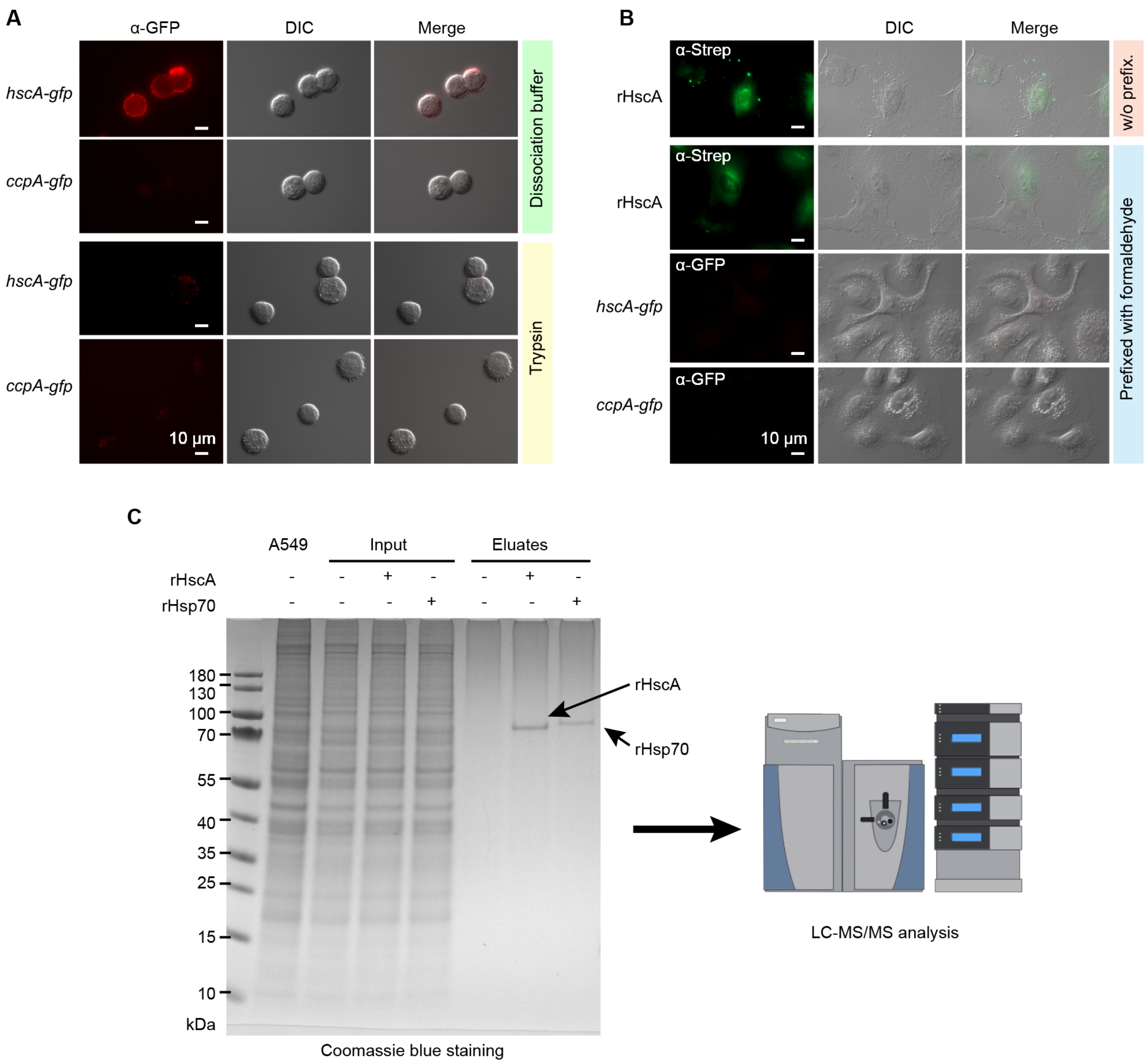

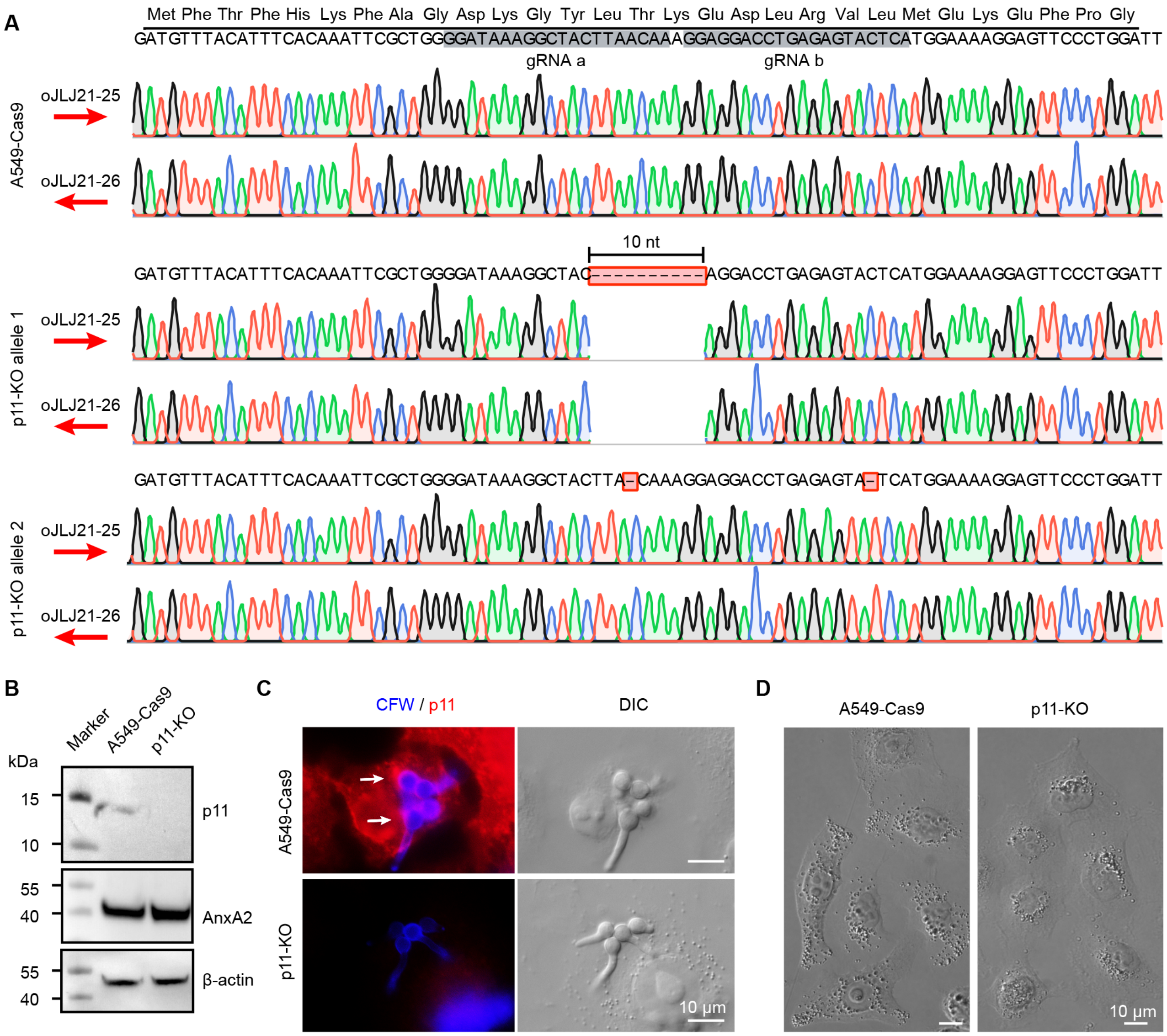

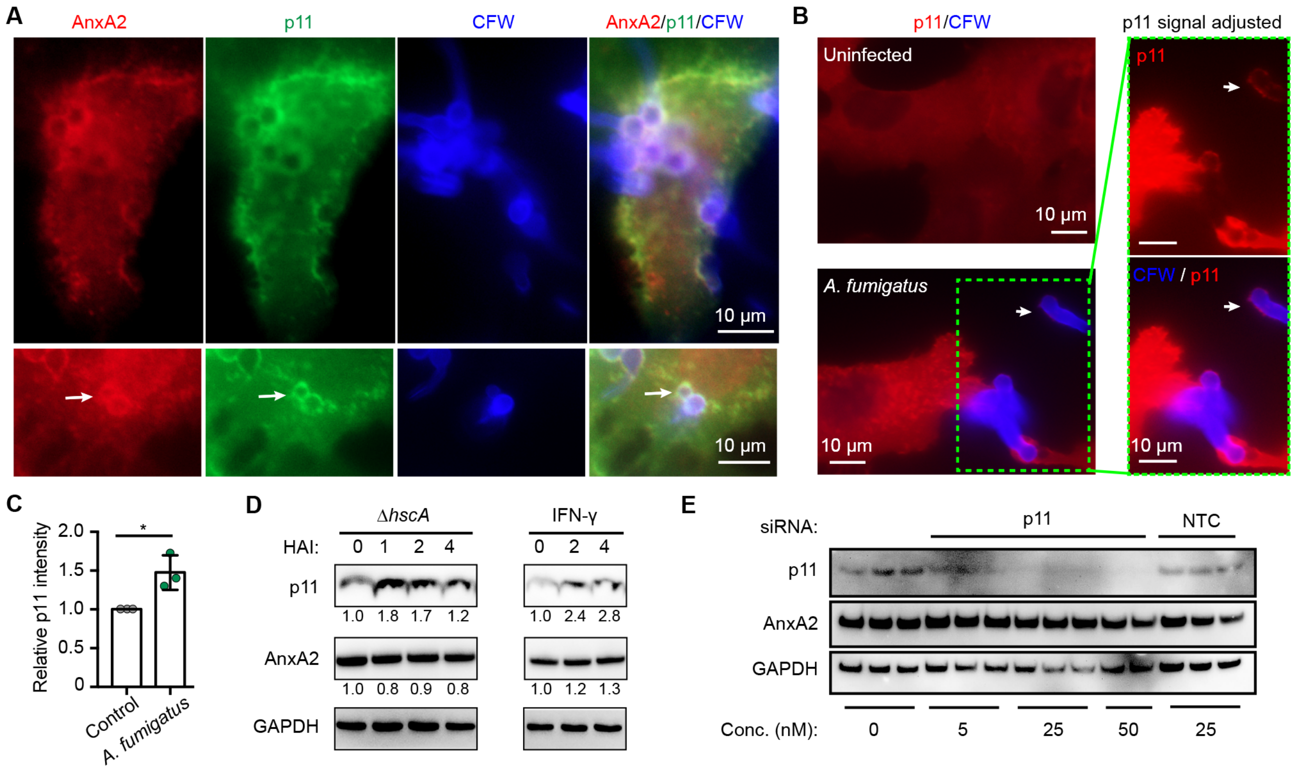

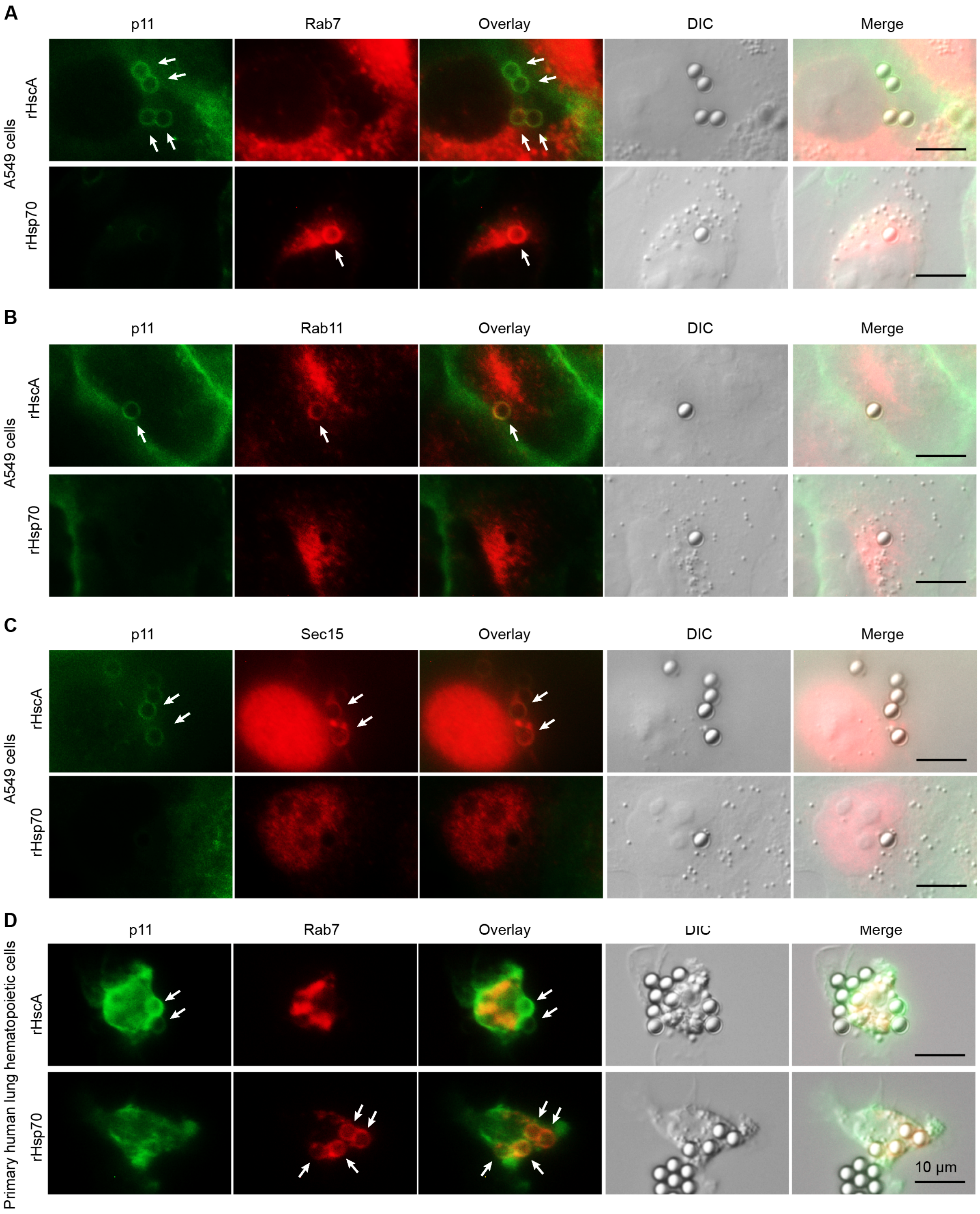

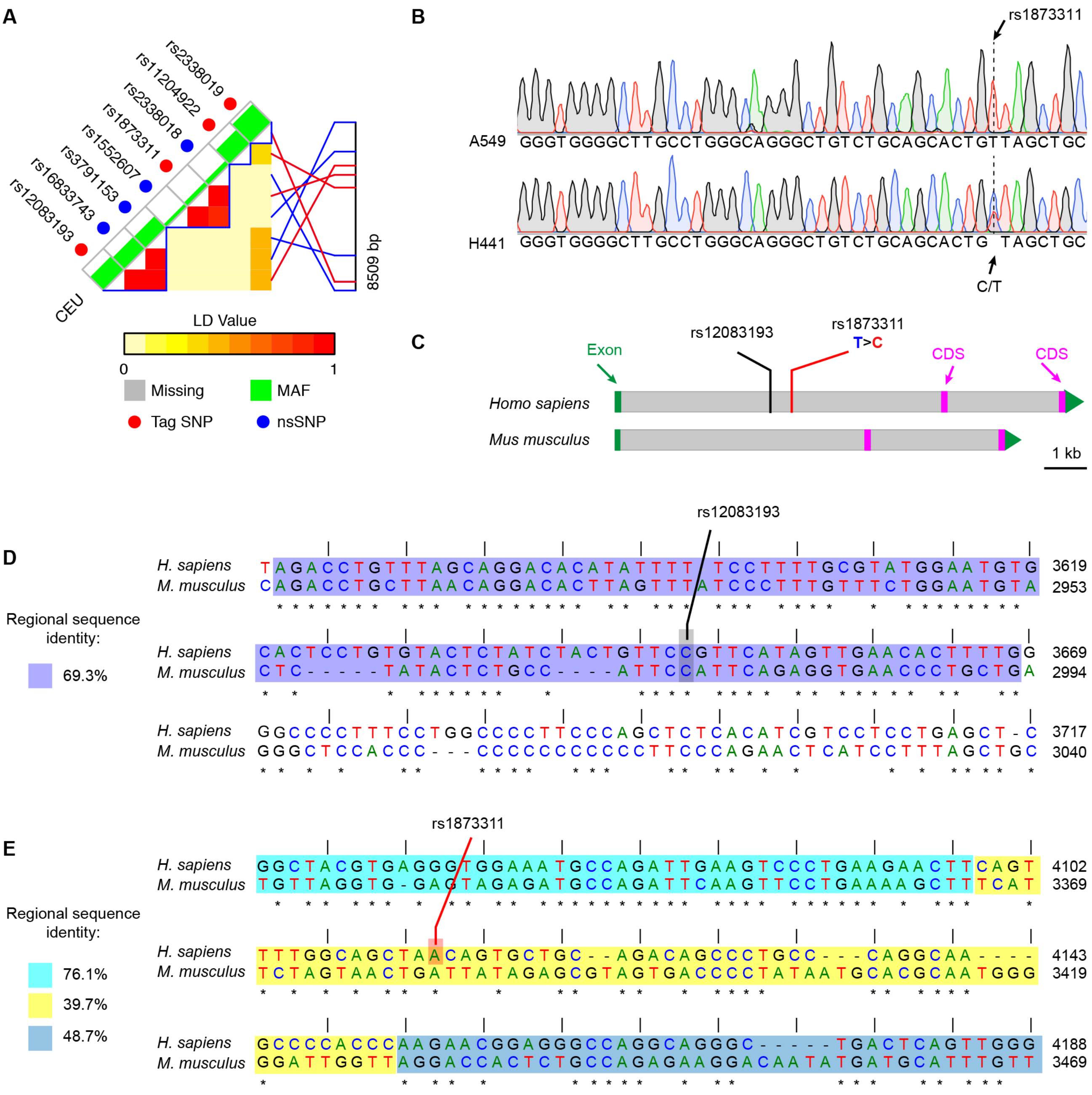

#### Fig. S7

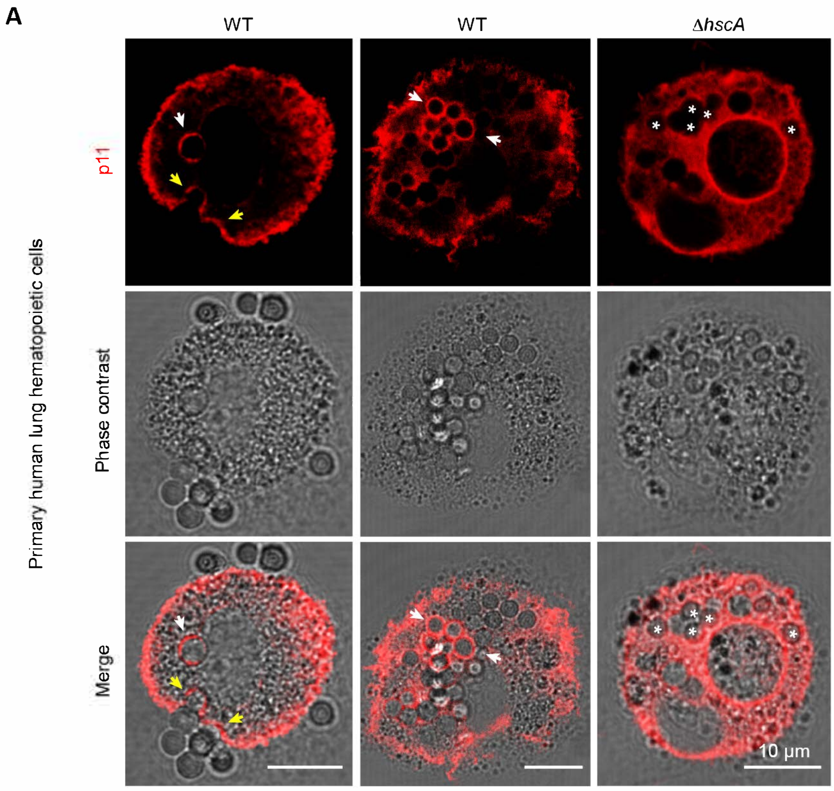


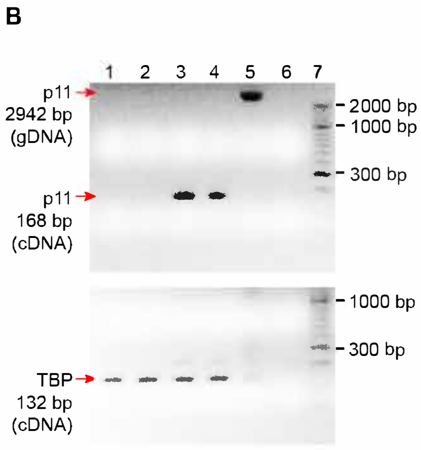


Lanes: 1: H441 uninfected (cDNA)
2: H441 infected with WT (cDNA)
3: A549 uninfected (cDNA)
4: A549 infected (cDNA)
5: H441 (gDNA)
6: $H_2O$

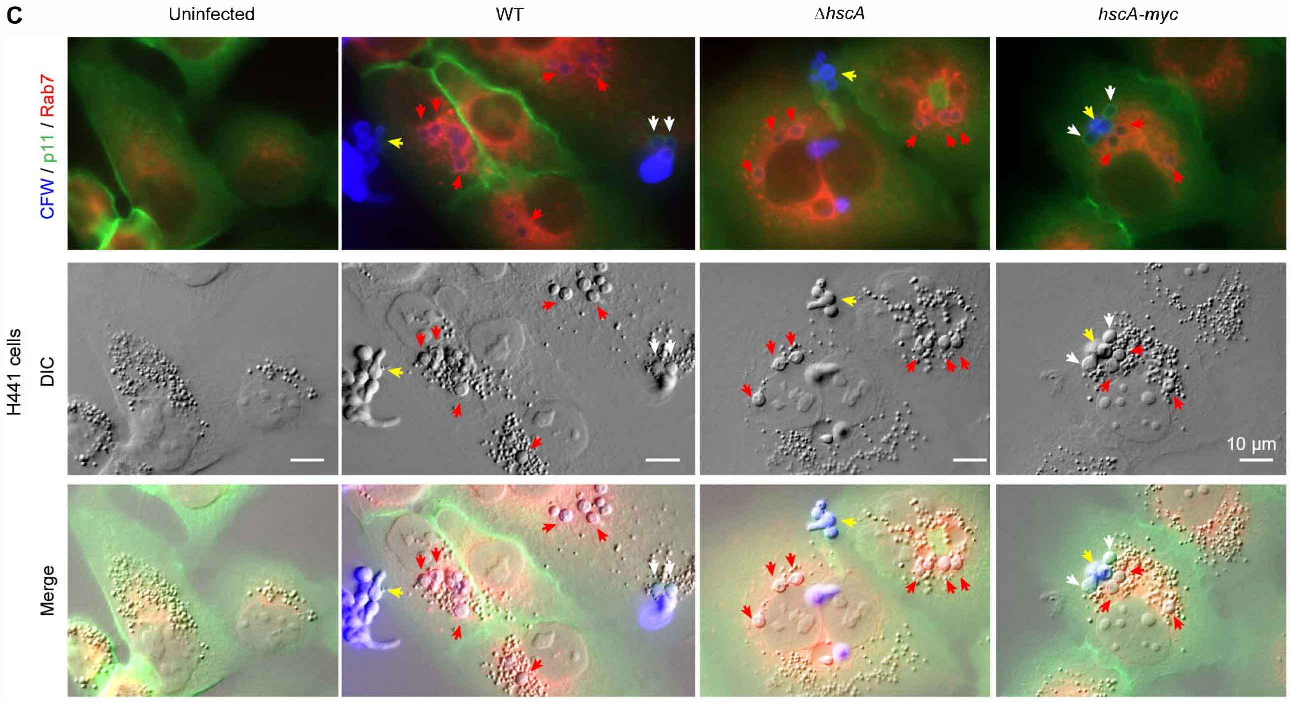
