## Supplementary Table S2 for "Re-direction of phagosomes to the recycling expulsion pathway by a fungal pathogen"

**Table S2. Frequency of *S100A10* genotypes among cases of IPA and controls, and association test results.**

| **SNP rs# (alleles)** | **Genome** | **Patients** | **Genotype, n (%)** | | | ***P* value** |
| --- | --- | --- | --- | --- | --- | --- |
|  |  |  | **A/A** | **A/a** | **a/a** |  |
| rs1873311  (T>C) | R | IPA | 94 (84.7) | 17 (15.3) | 0 (0.0) | 0.96 |
|  |  | Controls | 314 (84.4) | 56 (15.1) | 2 (0.5) |  |
|  | D | IPA | 98 (88.3) | 11 (9.9) | 2 (1.8) | 0.026 |
|  |  | Controls | 304 (81.7) | 67 (18.0) | 1 (0.3) |  |
| rs12083193  (G>C) | R | IPA | 55 (49.6) | 47 (42.3) | 9 (8.1) | 0.97 |
|  |  | Controls | 179 (48.1) | 162 (43.6) | 31 (8.3) |  |
|  | D | IPA | 54 (48.7) | 41 (36.9) | 16 (14.4) | 0.24 |
|  |  | Controls | 172 (46.2) | 164 (44.1) | 36 (9.7) |  |
| rs11204922  (T>C) | R | IPA | 38 (34.2) | 57 (51.4) | 16 (14.4) | 0.96 |
|  |  | Controls | 122 (32.3) | 196 (53.5) | 54 (14.2) |  |
|  | D | IPA | 42 (37.8) | 54 (48.7) | 15 (13.5) | 0.71 |
|  |  | Controls | 125 (33.6) | 192 (51.6) | 55 (14.8) |  |
| rs2338019  (T>C) | R | IPA | 33 (29.7) | 53 (47.8) | 25 (22.5) | 0.74 |
|  |  | Controls | 124 (33.3) | 173 (46.5) | 75 (20.2) |  |
|  | D | IPA | 30 (27.0) | 54 (48.7) | 27 (24.3) | 0.32 |
|  |  | Controls | 126 (33.9) | 173 (46.5) | 73 (19.6) |  |

SNP, single nucleotide polymorphism; IPA, invasive pulmonary aspergillosis; R, recipient; D, donor. The major and minor alleles are represented by the first and second nucleotides, respectively. A and a indicate distinct alleles of the same gene. *P* value is for Fisher’s exact *t* test.
