## Supplementary Table S3 for "Re-direction of phagosomes to the recycling expulsion pathway by a fungal pathogen"

**Table S3. Baseline characteristics of transplant recipients enrolled in the study.**

| **Variables** | **IPA**  **(n=111)** | **No IPA (n=372)** | **P value** |
| --- | --- | --- | --- |
| **Age at transplantation, no (%)** |  |  |  |
| ≤20 years | 16 (14.4) | 81 (21.8) | 0.150 |
| 21 – 40 years | 30 (27.0) | 108 (29.0) |  |
| >40 years | 65 (58.6) | 183 (49.2) |  |
| **Gender, no (%)** |  |  |  |
| Female | 48 (43.2) | 158 (42.5) | 0.870 |
| Male | 63 (56.8) | 214 (57.5) |  |
| **Underlying disease, no. (%)** |  |  |  |
| Acute leukemia | 61 (55.0) | 197 (53.0) | 0.223 |
| Chronic lymphoproliferative diseases | 16 (14.4) | 67 (18.0) |  |
| Myelodysplastic/myeloproliferative diseases | 17 (15.3) | 34 (9.1) |  |
| Chronic myeloproliferative diseases | 8 (7.2) | 21 (5.6) |  |
| Aplastic anemia | 6 (5.4) | 29 (7.8) |  |
| Other | 3 (2.7) | 24 (6.5) |  |
| **Transplantation type, no. (%)** |  |  |  |
| Matched, related | 36 (32.4) | 175 (47.0) | 0.009 |
| Matched, unrelated | 40 (36.0) | 91 (24.5) |  |
| Mismatched, related | 0 (0.0) | 8 (2.2) |  |
| Mismatched, unrelated | 35 (31.5) | 98 (26.3) |  |
| **Graft source, no. (%)** |  |  |  |
| Peripheral blood | 91 (82.0) | 306 (82.3) | 0.645 |
| Bone-marrow | 19 (17.1) | 57 (15.3) |  |
| Cord blood | 1 (0.9) | 9 (2.4) |  |
| **Disease stage, no. (%)** |  |  |  |
| First complete remission | 59 (53.2) | 204 (54.8) | 0.940 |
| Second or subsequent remission, or relapse | 19 (17.1) | 63 (17.0) |  |
| Active disease | 33 (29.7) | 105 (28.2) |  |
| **Conditioning regimen, no (%)** |  |  |  |
| RIC | 79 (71.2) | 250 (67.2) | 0.452 |
| Myeloablative | 32 (28.8) | 122 (32.8) |  |
| **CMV serostatus of donor and recipient, no. (%)** |  |  |  |
| D-/R+ or D+/R+ | 94 (84.7) | 331 (89.0) | 0.214 |
| D-/R- or D+/R- | 17 (15.3) | 41 (11.0) |  |
| **Duration of neutropenia, mean days (range)†** | 13.2 (8 – 39) | 14.0 (5 – 35) | 0.504 |
| **Acute GVHD, no. (%)** |  |  |  |
| No GVHD or grades I – II | 77 (69.4) | 325 (87.4) | <0.001 |
| Grades III – IV | 34 (30.6) | 47 (12.6) |  |
| **Antifungal prophylaxis, no. (%)‡** |  |  |  |
| Fluconazole | 55 (49.6) | 149 (40.1) | 0.036 |
| Posaconazole | 31 (27.9) | 120 (32.3) |  |
| Other | 9 (8.1) | 15 (4.0) |  |
| None or unknown | 16 (14.4) | 88 (23.7) |  |

Chronic lymphoproliferative diseases included cases of chronic lymphocytic leukemia, multiple myeloma, and B- and T-cell lymphomas. Chronic myeloproliferative diseases included cases of chronic myelogenous leukemia and primary myelofibrosis. Other diseases included cases of idiopathic medullar aplasia, lymphohistiocytosis, hemoglobinopathies and paroxysmal nocturnal hemoglobinuria. RIC, reduced intensity conditioning; CMV, cytomegalovirus; D, donor; R, recipient; GVHD, graft-versus-host-disease. †Neutropenia was defined as ≤0.5×10^9^ cells/L. ‡Other antifungals used in prophylaxis included voriconazole, liposomal amphotericin B, itraconazole and caspofungin. P values were calculated by Fisher’s exact probability t-test or Student’s t-test for continuous variables.
